# Improved detection of differentially abundant proteins through FDR-control of peptide-identity-propagation

**DOI:** 10.1101/2024.11.15.623880

**Authors:** Alexander J. Solivais, Hannah Boekweg, Lloyd M. Smith, William S. Noble, Michael R. Shortreed, Samuel H. Payne, Uri Keich

## Abstract

The goal of proteomics is to identify and quantify peptides and proteins within a biological sample. Almost all algorithms for the identification of peptides in LC-MS/MS data employ two steps: peptide/spectrum matching and peptide-identity-propagation (PIP), also known as match-between-runs. PIP was originally envisioned as a backup method to overcome measurement stochasticity. However, current PIP implementations can routinely account for up to 40% of all results, with that proportion rising as high as 75% in single-cell proteomics. Unlike peptide identities derived through peptide/spectrum matches, for which error estimation has been strictly enforced for decades, peptide identities derived through PIP have not historically been subject to statistical evaluation. As an indispensable component of label free quantification, PIP needs a simple and statistically rigorous method for estimating its error rates. Although several tools claim to control the false discovery rate (FDR) of PIP, these claims cannot be validated as there is currently no accepted method to assess the accuracy of the stated FDR. We present a method for FDR control of PIP, called PIP-ECHO, and devise a rigorous protocol for evaluating FDR control of any PIP method. Using three different benchmark datasets, we evaluate PIP-ECHO alongside the PIP procedures implemented by FlashLFQ, IonQuant, and MaxQuant. These analyses show that only PIP-ECHO can accurately control the FDR of PIP at 1% across all datasets, including single cell. When analyzing spike-in datasets where different known amounts of yeast or E. coli peptides are added to a constant background of human peptides, PIP-ECHO increases both the accuracy and sensitivity of differential expression analysis, yielding 53% more differentially abundant proteins than MaxQuant and 146% more than IonQuant.

## Introduction

Liquid chromatography-tandem mass spectrometry (LC-MS/MS) is the basis of modern proteomics. In the most common form of proteomic analysis, bottom-up proteomics, complex mixtures of peptides obtained through the enzymatic digestion of proteins are separated using LC and analyzed using MS/MS. The mass spectrometer records MS1 scans, which contain the mass-to-charge ratios (*m/z* ‘s) of all analytes eluting at a given time. In data-dependent acquisition (DDA), a common MS/MS analysis method, each MS1 spectrum is followed by a few MS2 fragmentation scans that are generated by isolating and fragmenting all analytes in a narrow *m/z* range corresponding to a series of isotopic peaks observed in the MS1 scan. Database search engines analyze the resulting data by matching the MS2 fragmentation spectra to theoretical fragmentation spectra generated using a peptide database. This procedure yields a list of peptide-spectrum-matches, or PSMs. We refer to peptides identified in this way as *peptide detections*.

A key component of the database search strategy is a statistical procedure to control the rate of errors among the reported PSMs. Some of the PSMs output by the search engine are correct, meaning that the matched peptide was present in the sample and generated the MS2 spectrum. Conversely, other PSMs are incorrect, meaning that the match was made by chance, and the reported peptide did not generate the MS2 spectrum. Canonically, target-decoy competition (TDC) is used to determine which PSMs are reported. TDC involves augmenting the peptide database with decoy peptides that are created by reversing or shuffling the original, or target, peptides. Because the decoy peptides are artificial, any PSM where a spectrum is matched to a decoy peptide is incorrect. Hence, the number of decoy peptides that are detected (in one or more PSMs) can be used to estimate and subsequently control the false discovery rate (FDR). The FDR is the expected value of the proportion of incorrect peptides among the reported target peptides, i.e., the false discovery proportion (FDP) [5, 8].

With the MS2-based peptide detection phase completed, we are often interested in quantifying the relative abundances of the detected peptides. Label-free quantification (LFQ) is a popular approach that uses information in MS1 scans to accomplish this task. Briefly, consider a peptide that was detected in a specific MS2 scan. As that peptide eluted it would have left a so-called “peak trace” across a short sequence of MS1 scans that are temporally adjacent to the detecting MS2 scan. These MS1 scans would contain the characteristic collection of *m/z* peaks corresponding to different isotopologues of the peptide (a graphical depiction of a “peak trace” is shown in Figure 1A, and a more thorough definition is provided in Methods). Pairing the detected peptide with its corresponding peak trace allows us to deduce the peptide’s relative abundance from the intensities of the peaks in the trace.

**Figure 1:**
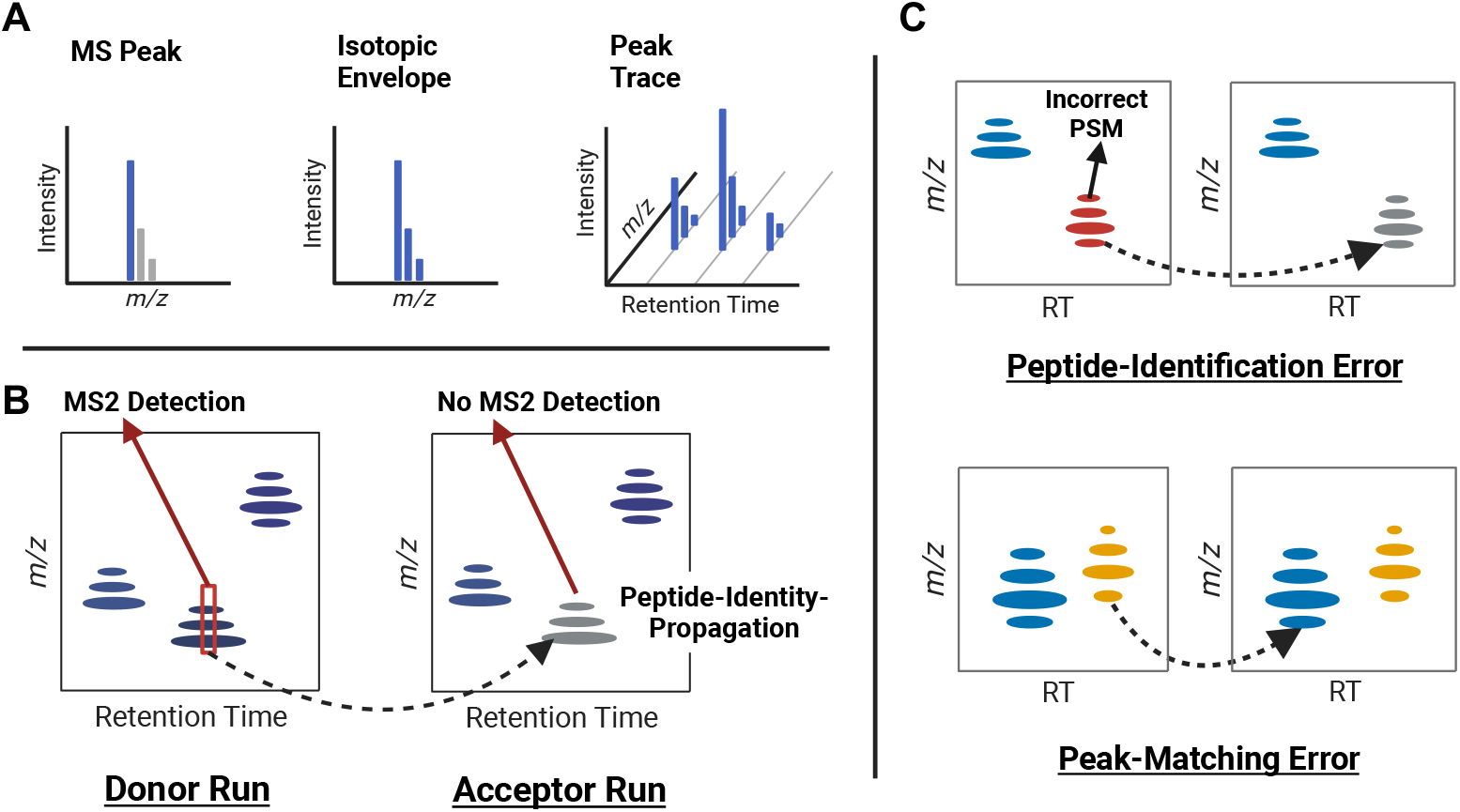
Key Terms: **A** Graphical depictions of MS1 spectra highlighting an MS peak, an isotopic envelope, and a precursor peak. **B** Illustration of the basic PIP workflow, where the precursor peak corresponding to an MS2-detection in the donor run is used to identify a matching precursor peak in the acceptor run, resulting in an PIP-detection. **C** Illustration of the two different types of error that can occur during PIP.

LFQ allows researchers to compare relative peptide or protein abundances between samples. Of course, we can only compare abundances of peptides that are consistently detected across multiple runs. However, inherent run-to-run variability is created by both real differences between biological samples and also technical instrument limitations. For example, a peptide may be present in multiple runs but only fragmented in some runs, resulting in missing peptide detections. For this reason, LFQ proteomics experiments yield quantitative data with 20–50% data incompleteness [24]. To mitigate this stochastic variability and improve data completeness, a family of algorithms has been developed to transfer the identity of detected peptides between runs based on pairings of peak traces. This procedure is known as “peptide-identity-propagation” (PIP) or sometimes “match-between-runs” (MBR).

The key idea in PIP is to identify pairs of peak traces with similar retention times and m/z values that occur in different LC-MS/MS runs. Only one of the two peak traces, the “donor,” is linked to an MS2 peptide detection. The other peak trace, the “acceptor”, originates in a run where that peptide was not MS2 detected. The identity of the peptide is then transferred from the donor to the acceptor peak. An illustration of the PIP procedure is shown in Figure 1B. This approach was originally designed to work with home-built ultra-high resolution FTICRs which lacked MS2 capability [13] but has since been adapted to many applications in proteomics [17, 4, 9]. In most current applications, PIP is thought of as a backup method to supplement peptide detections from MS2 spectra. In practice, PIP can be responsible for a large fraction of all identified peptides, i.e., peptides that are paired with specific MS1 features. In single-cell proteomics in particular, PIP can account for more than 75% of all peptide identifications [26].

Because PIP generates a large fraction of all peptide identifications, it is essential that we understand and control errors in the method. Similar to the PSMs produced by database search engines, many of the peptide identifications arising from PIP are correct: the MS1 peaks in the acceptor run were generated by the same peptide that was MS2-detected and linked to the donor peak trace. However, others are not. Thus, as a vital component of quantitative proteomics, it is essential that we develop procedures for controlling the error rate of PIP.

Error analysis in PIP is complex, as two distinct types of errors can occur. First, *peak-matching errors* occur when the donor-acceptor peak traces are incorrectly paired, either because the acceptor peak trace was generated by a different analyte or is a noise-generated artifact. Second, *peptide-identification errors* occur when the identity of the donor peptide that was propagated was falsely matched to the donor peak trace prior to PIP. This type of error mostly occurs when the donor peak trace is linked to an incorrect peptide detection. If the MS2 detection procedure used an FDR threshold of 1%, then on average we expect that approximately 1% of all peptides used as donors in the PIP process will be incorrect, resulting in peptide-identification errors. Both types of errors are depicted in Figure 1C. Although this type of error is not caused by failures of the PIP algorithm, these errors do have important implications for the accuracy of the results. When a peptide-identification error occurs, even if the peak traces in the donor and acceptor are correctly matched, they now have an incorrect label and contribute to the overall error rate among reported transfers.

Initial implementations of PIP did not provide any measure of statistical confidence. Subsequent work evaluated the potential for errors in the method but did not provide a general solution for how to estimate statistical confidence [1]. For example, with a two-proteome experimental design, Lim et al. used a combined human+yeast dataset as donors and human-only data as acceptors to quantify error rates [12]. In this approach, some of the peak-matching errors can easily be detected; in particular, any yeast peptide matched in a human-only sample constitutes such an error. Surprisingly, 44% of detected yeast proteins were incorrectly transferred to a human-only sample. Although most of these false transfers were “one-hit-wonders,” meaning that only one peptide from the protein was transferred, this demonstrates that false transfers can have an outsized impact on errors when rolled up to the protein level. This is especially concerning in single cell proteomics, where protein detections frequently rely on only one peptide.

More recently, several methods have been introduced that attempt to account for peak-matching errors during PIP. For example, IonQuant [6] attempts to sample from the null distribution of incorrect feature– feature matches by looking for matches to “decoy” features in the acceptor run. These decoy features’ masses are selected to be shifted by 5–11 m/z from the donors’ feature masses; hence, they presumably represent incorrect transfers. Quandenser [20] uses a similar strategy to model the null distribution, relying on decoy acceptor features that are offset by 5 m/z from the donors’ masses. However, neither tool accounts for possible peptide-identification errors, making the implicit assumption that all donor peptide detections are correct. While the combination of Quandenser with triqler [21] does consider both types of error, this is done internally as part of the overall goal of producing a list of differentially abundant proteins, and it is not clear how this method can be used for controlling the error in the PIP process itself.

Here we introduce PIP Error Control via Hybrid cOmpetition (PIP-ECHO), a procedure to carry out PIP while accounting for both peak-matching and peak-identification errors. As its name suggests, PIP-ECHO leverages two types of competition to control the FDR among the reported list of transferred peptide identities. The first competition mimics the canonical target-decoy peptide competition used in FDR control for database search. The second competition involves generating matches to peak traces detected at randomized retention times. This step is similar in spirit to the methods employed by Quandenser and IonQuant. The heart of PIP-ECHO is the novel way in which it leverages both types of competitions to estimate and control the overall FDR for PIP.

Although PIP-ECHO is analytically motivated, it is important to gauge how successful the method is in controlling the FDR in practice. This goal leads to the second major contribution of this paper, a procedure that uses the two-proteome experimental design to quantifiably gauge how well a PIP procedure controls the overall FDR among its reported list of transfers. Applying this novel procedure to multiple two-proteome datasets, including one simulating single-cell proteomics data, we show that MaxQuant, IonQuant, and the previous version of FlashLFQ all fail to control the overall FDR for PIP. Moreover, these experiments show that the new version of FlashLFQ, which implements PIP-ECHO, successfully controls the overall PIP FDR while delivering a comparable number of discoveries as MaxQuant and IonQuant.

We demonstrate the practical utility of PIP-ECHO by performing an experiment designed to detect proteins that are differentially abundant in two conditions. Specifically, we used FlashLFQ+PIP-ECHO, as well as MaxQuant and IonQuant, to infer a PIP-boosted list of MS1-quantified peptides from a spike-in dataset originally published by Shen et al. [15]. This dataset was generated by varying the proportion of *E. coli* proteins that were spiked into an essentially fixed concentration of human proteins. Thus, the *E. coli* peptides are differentially abundant with known expected fold-change values, whereas the human peptide abundances are not expected to change. We subsequently applied two types of differential abundance analyses comparing the empirical fold change computed from the MS1-quantified peptides with the expected one. In both analyses, we found that when the error rate is held constant, FlashLFQ+PIP-ECHO allows us to identify substantially more peptides and proteins at the expected fold change than either MaxQuant or IonQuant. In the second analysis, the results from FlashLFQ+PIP-ECHO enabled the discovery of 53% and 146% more differentially abundant proteins as compared to MaxQuant and IonQuant, respectively.

PIP-ECHO is implemented in a new version of FlashLFQ, which is open source and freely available both as a stand-alone program and as a component of the MetaMorpheus software suite [19].

## Results

### PIP-ECHO: controlling for both types of errors

PIP-ECHO is designed to report as many correct PIPs as possible while controlling the overall FDR. This requires a procedure to estimate the number of incorrect PIPs included in a set of candidate PIPs, as well as a scoring function that quantifies the quality of each putative PIP. To estimate the number of incorrect PIPs, we need to account for both peak-matching and peptide-identification errors. PIP-ECHO uses two types of competitions to estimate, and therefore control, the frequency of these two types of errors.

Peptide-identification errors are accounted for using the first competition, which relies on the canonical TDC conducted during the database search step, where each MS2 spectrum was searched against a concatenated target-decoy database. PIP-ECHO transfers the identity of both target and decoy peptides that were detected (at a given FDR threshold) during this database search step (Figure 2A). This produces target-peptides PIPs, where the identity of a target peptide is transferred, and decoy-peptide PIPs, where the identity of a decoy peptide is transferred. By counting the number of decoy-peptide PIPs, we are able to estimate the number of PIPs where the identity of an incorrectly detected target peptide is transferred, i.e., peptide-identification errors.

**Figure 2:**
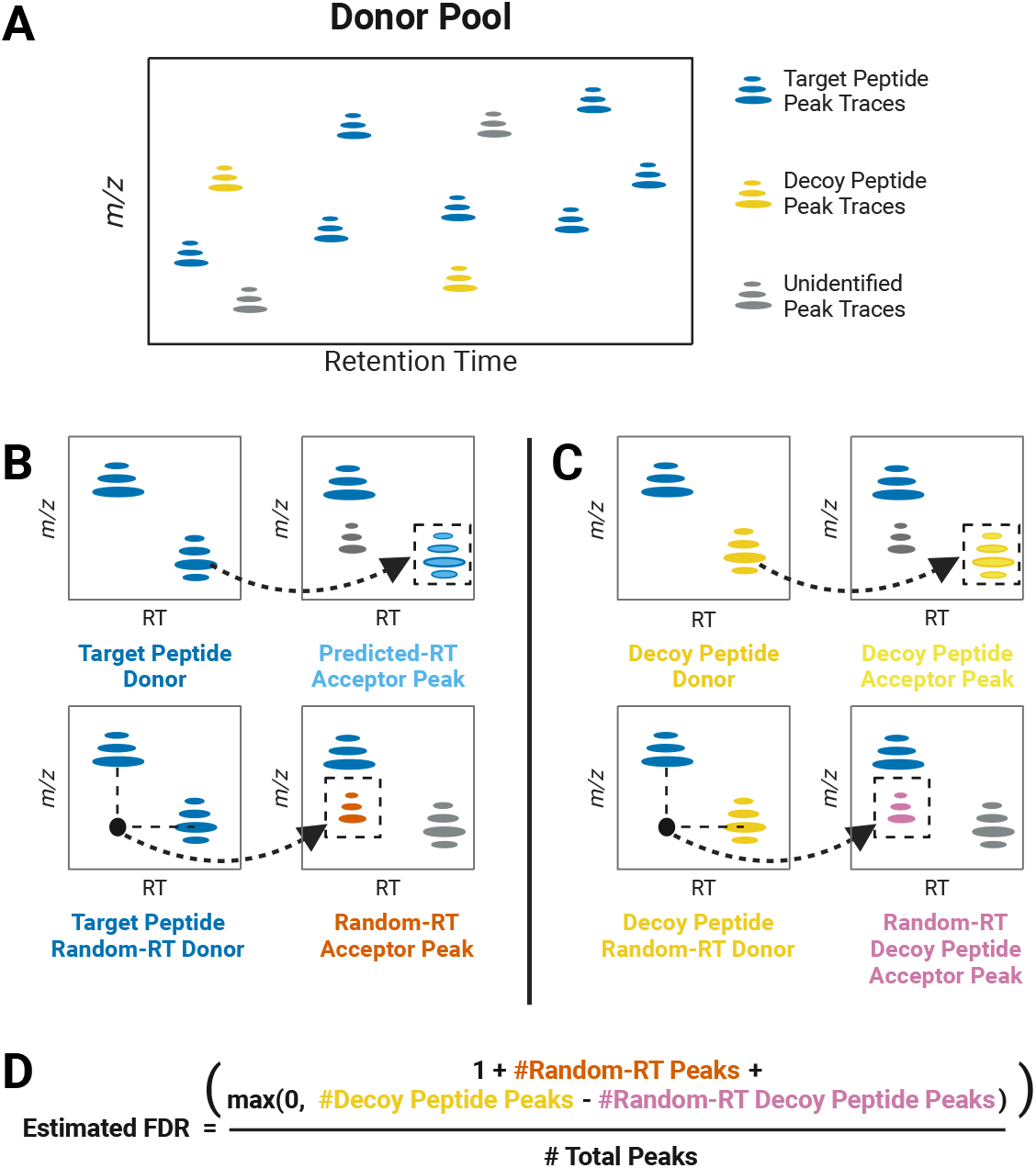
Workflow for PIP-ECHO FDR Control: **A** A depiction of the MS1 signal from a single LC-MS run showing peak traces associated with target and decoy peptides. **B** Illustrated example of predicted and random-RT peak selection. **C** Illustrated example of decoy peptide peak selection and decoy peptide, random-RT peak selection. **D** The equation used to calculate the estimated FDR in PIP-ECHO.

Peak-matching errors are accounted for using the second competition, where PIPs that are located at a predicted retention time (RT) — the retention time of the donor peptide mapped onto the acceptor run — are competed against PIPs located at a randomized RT. These random RTs are selected by considering all peptides in the donor run whose mass is “close but not too close” to that of the donor peptide (Figure 2B-C). Then, one of these peptides is selected at random, and its RT is mapped onto the acceptor run. Finally, we attempt to locate a peak trace with the same mass and isotopic distribution as the donor peptide but anchored at this randomized RT. Such random-RT peak traces model peak-matching errors: the peak trace is observed at the wrong RT and therefore should not be linked to the donor peak trace.

Assuming that an incorrect PSM is equally likely to involve a target or a decoy peptide, as is canonically done in TDC, the number of decoy-peptide PIPs allows us to estimate the number of incorrect target-peptide PIPs. We similarly assume that an incorrect transfer for a given donor peak trace is equally likely to involve the predicted RT as it is to involve the random RT. Accordingly, PIP-ECHO competes each such pair of predicted-RT and random-RT transfers, keeping only the higher scoring of the two, and uses the number of times a random-RT PIP wins the competition to estimate the number of target-peptide PIPs that are actually peak-matching errors.

PIP-ECHO combines the information captured by these two competitions to estimate and control the FDR in each acceptor run separately. For each run, let 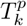 be the number of candidate target-peptides PIPs with a predicted RT among the top *k* scoring candidates, and let 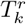 be the corresponding number of target-peptide PIPs with a randomized RT among the same top *k*. Similarly, let 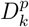 and 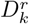 be the analogous numbers of decoy-peptide PIPs. The assumptions above imply that 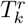 is an unbiased estimate of the number of peak-matching errors. Similarly, 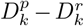 is an estimate of the number of peptide-identification errors (with correctly matched peaks). 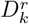 is subtracted from 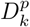 to avoid double-counting of errors, as incorrect peak transfers are accounted for in the first term. Because 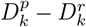 can be negative, we take the maximum of it and 0, arriving at the following estimate for the FDR among the 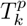 top scoring predicted-RT target-peptide PIPs:

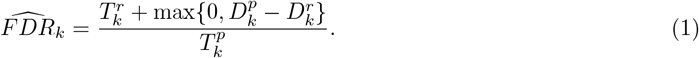

In attempting to control the FDR among the run’s reported PIPs at level *α*, PIP-ECHO then utilizes the above estimate analogously to [8]: by adding +1 to the numerator and looking for the maximal *k* such that the estimate is still ≤ *α*:

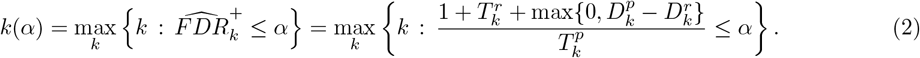

The procedure then reports the corresponding 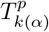 top scoring predicted-RT-based target PIPs.

In order to maximize PIP-ECHO’s power, i.e., the number of PIPs reported at a given FDR threshold, we need a scoring function that is able to distinguish between correct and incorrect transfers. We therefore developed an iterative rescoring that uses gradient boosted decision trees to gradually learn to separate correct and incorrect transfers, modeled by predicted-RT and random-RT PIPs, respectively. Like Percolator [10], we use cross-validation to ensure that the tree learns to distinguish between correct and incorrect transfers rather than only learning how to distinguish between predicted and random-RT PIPs (i.e., overfitting).

### Estimating false discovery proportions in PIP using two-proteome datasets

A procedure that controls the FDR in practice controls only the expected value of the false discovery proportion (FDP). The latter varies from dataset to dataset but should on average be ≤ *α*, where *α* is the procedure’s target FDR threshold. For such a procedure, we expect that when the number of discoveries is large, the actual FDP should be at most about *α*. In our case, the FDP is the proportion of PIPs in the final list of reported PIPs that involve either a peak-matching or a peptide-identification error.

Computational entrapment procedures are often used to assess the FDR control of database search tools by estimating the FDP in the reported result [25]. However, the two types of errors present in the PIP context require a different experimental design. We therefore present a new entrapment procedure to estimate the FDP of PIP using data from two-proteome experiments [12]. In a two-proteome experiment, cell lysate digests from two different species are used to generate two samples: a pure sample, which contains peptides only from species A; and a mixed sample, which contains a mix of peptides from both species A and species B. Each sample is analyzed by LC-MS/MS multiple times to generate replicates, and these replicates are analyzed together.

Our procedure focuses on estimating the FDP due to peptide-identification and peak-matching errors among all PIPs reported in the pure (species A) runs. Analogously to the canonical entrapment setup we estimate the number of peptide-identification errors by adding entrapment peptides that are not expected to be in the sample to the target database. Any PIP that is derived from an entrapment donor peptide is therefore presumed to be a peptide-identification error, and similarly to TDC we can use those observed errors to estimate the number of unobserved peptide-identification errors among the target-peptide PIPs.

To estimate the number of peak-matching errors, we note that each can be classified either as a native-peak error, where the donor peptide is present in the acceptor run but it is matched to an incorrect peak trace, or as a foreign-peak error, where the donor peptide is not present in (or foreign to) the acceptor run. In a two-proteome experiment, the pure samples only contain peptides from species A. Therefore, any PIP reported in a pure sample with a donor peptide from species B clearly constitutes an observed foreign peak-matching error. Moreover, we assume that the sample from species A is shared across all runs and hence that there are no other PIPs with foreign-peak errors in the pure runs (entrapment PIPs are already counted as peptide identification errors).

In order to estimate the number of native-peak errors that occur in the pure sample runs, we mask a subset of peptide detections by editing the data to remove MS2 spectra in which the peptides were detected. We then reanalyze the edited data, relying on PIP to locate peak traces corresponding to the masked peptides. The peak traces identified through PIP in the reanalysis are compared to the peak traces identified based on MS2 spectra in the first analysis. If the two peak traces have apex retention times separated by more than 1% the length of the LC gradient) (54 seconds for a 90 minute gradient), it is considered a native peak error.

Finally, we combine the estimated number of native and foreign peak-matching errors with the estimated number of peptide-identifications errors to estimate the overall PIP FDP. This approach enables an accurate estimation of the FDP of PIP during analysis of two-proteome datasets (see Methods for more details).

### Only PIP-ECHO consistently controls the overall FDR

We compared the FDP among the PIPs reported by PIP-ECHO, MaxQuant [3], IonQuant [6], and FlashLFQ[14] by applying the above method to three different two-proteome datasets. The first dataset, originally published by Lim et al., consists of mixed samples containing human and yeast peptides and pure samples containing only human peptides. The second dataset consists of mixed samples containing human and *E. coli* peptides, and pure samples containing only human peptides. The final dataset, originally published by Truong et al., was designed to mimic a single-cell proteomics experiment, where it is common to analyze single-cell samples alongside samples derived from bulk cell lysate in order to increase the number of peptides detected in single cells [22]. Specifically, ten nanograms of peptides were injected for each mixed human and yeast sample replicate and only 200 picograms of peptides (equivalent to the expected amount in a single human cell) were injected for each pure human sample replicate.

The resulting FDP analysis suggests that only PIP-ECHO appears to control the FDR in the single-cell equivalent dataset (Figure 3): at the 1% PIP FDR threshold PIP-ECHO’s estimated FDP is 1.0% compared with IonQuant’s 3.4%. For FlashLFQ v1.0 we could only filter at the donor level (PSMs were filtered to an FDR of 1%), the same applies to MaxQuant (PSM and protein level FDR were both set to 1%) and the corresponding estimated PIP FDPs were 6.1% and 2.0%, respectively. These results suggest that existing PIP tools are prone to errors when applied to single-cell proteomics data.

**Figure 3:**
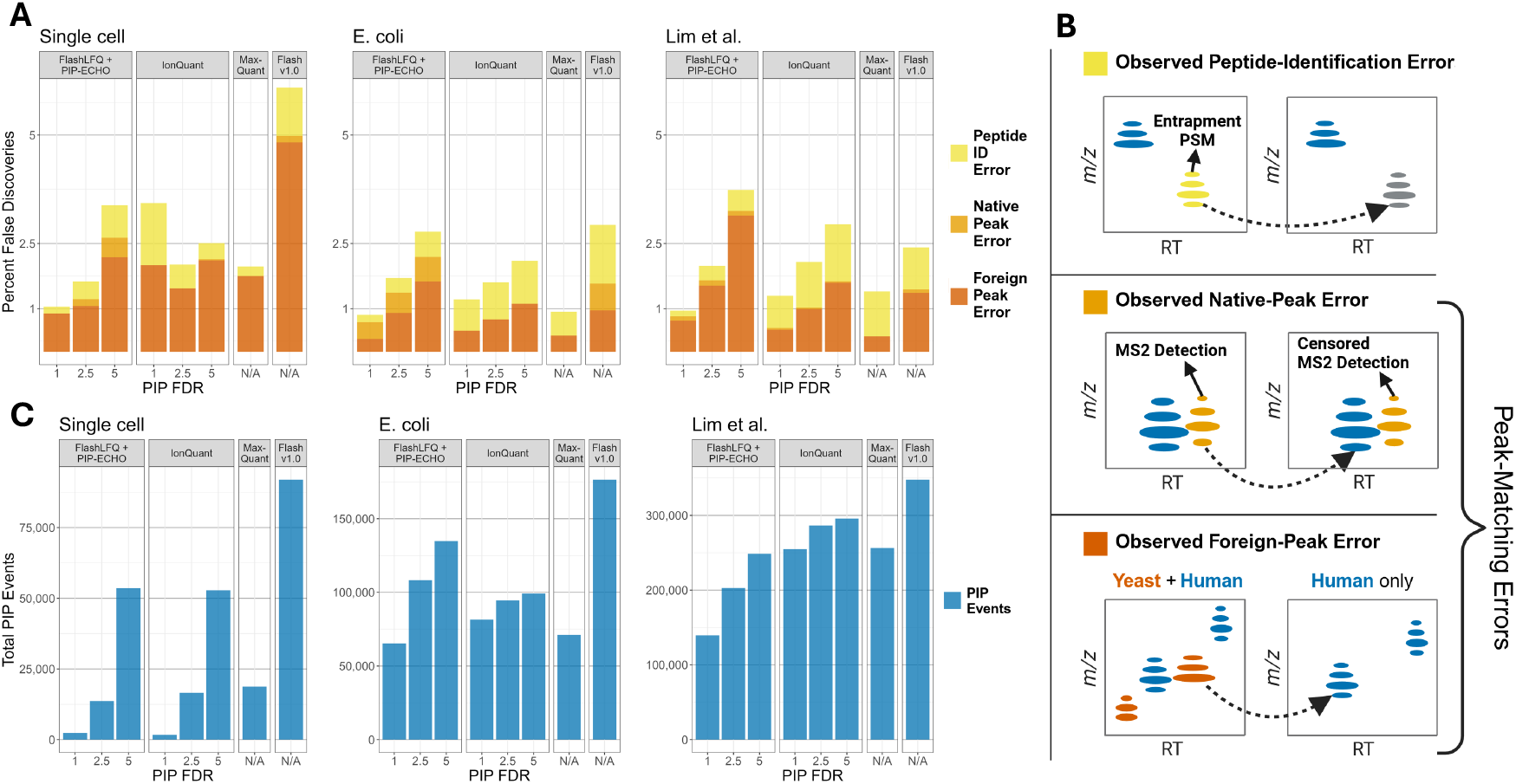
PIP Errors by PIP-ECHO (as implemented in FlashLFQ v2.0), IonQuant, MaxQuant, and FlashLFQ (v1.0): **A** Plot showing the number of PIP detection errors made by each software tool when evaluated using each of three different two-proteome datasets. For IonQuant and PIP-ECHO, three different PIP FDR thresholds were tested: 1, 2.5, and 5%. For FlashLFQ v1.01 and MaxQuant, N/A denotes that the software reports a list of PIP discoveries without controlling the FDR. **B** Illustration of how we estimate the three different types of errors that can occur during PIP. **C** Plot showing the number of PIPs reported by each software tool.

For the other two datasets, FlashLFQ’s estimated FDP is about 2.5% for both, while the estimated FDP for MaxQuant is 1.4% and 0.9%. While IonQuant reports a higher number of discoveries than PIP-ECHO at the same 1% threshold, the estimated FDP is higher than 1%: 1.3% and 1.2%, whereas PIP-ECHO’s estimated FDP is consistently at or below 1%.

Looking closer at the breakdown of the estimated FDP we find that for MaxQuant, IonQuant, and FlashLFQ v1.0, approximately half of the errors among PIPs reported for the *E. coli* and Lim et al. datasets are peptide-identification errors — errors that these tools do not account for. The corresponding estimated fraction of peptide-identification errors in PIP-ECHO’s case is increasing with the overall FDR threshold, but remains significantly lower. This is thanks to a heuristic that PIP-ECHO employs, where it sets the donor peptide FDR threshold at one fifth of the overall PIP FDR threshold, e.g., for a PIP-FDR of 1%, the considered donor peptides will be those that are discovered in a database-search using an FDR cutoff of 0.002 (0.2%). Note that this heuristic should only impact the sensitivity of PIP-ECHO and has no effect on its FDR control. In Figure S1 we show a similar breakdown of PIP-ECHO’s estimated FDP where we disable this heuristic, defining the list of potential donor peptides using a fixed 0.2% database-search FDR cutoff, and consider a continuous range of FDR thresholds.

### FlashLFQ+PIP-ECHO quantification improves downstream differential abundance analysis

We next compared how the quantitative results generated by IonQuant, MaxQuant, FlashLFQ v1.0, and FlashLFQ+PIP-ECHO impact downstream differential abundance analysis. In order to do so, we analyzed a spike-in dataset originally published by Shen et al. [15]. This dataset consists of *E. coli* and human lysate digests mixed at five different ratios, ranging from 3% to 9% total *E. coli* lysate (wt/wt). Four technical replicates were used for each concentration, resulting in 20 LC-MS/MS runs in total. Pairwise analyses were performed between each of the five concentrations for a total of 10 pairwise analyses. Notably, because this is a controlled dataset we know which peptides and proteins are differentially expressed and what is the expected log-fold change (*E. coli* ‘s), and which are not supposed to be changed (human). This setup allows us to compute the FDP in any list of presumed differentially expressed analytes.

A common approach to differential expression analysis is to apply *limma*, a popular t-test based `R` package. Originally designed for the analysis of microarray data, *limma* was adopted by the proteomics community to detect peptides or proteins that are differentially expressed between alternate experimental conditions [18]. We used *limma* to analyze the outputs from IonQuant, FlashLFQ v1.0, and FlashLFQ+PIP-ECHO and find that the latter delivers consistently more discoveries at comparable FDP levels for every selected level of PIP FDR (including no -PIP and all PIPs). For example, using FlashLFQ+PIP-ECHO, *limma* is able to detect, at the 1% DE FDR level, 22% more proteins than when using IonQuant, while still yielding a DE FDP of less than 1%, and this holds for all PIP FDR thresholds (Figure S2A). Unsurprisingly, when PIP is disabled, *limma*’s analysis based on FlashLFQ v1.0 delivers similar performance to that based on FlashLFQ+PIP-ECHO (Figure S2A-D). However, when PIP is enabled (FlashLFQ v1.0 does not allow PIP FDR control) we see that *limma*’s analysis based on FlashLFQ v1.0 often significantly exceeds the selected FDR threshold (panels A & C) and is consistently substantially higher than the FDP of the corresponding analysis based on FlashLFQ+PIP-ECHO even when the latter uses the 100% FDR threshold. Presumably, this is thanks to PIP-ECHO’s improved ability to select correct transfers, something we will return to below.

Although the *limma* package is popular and commonly used for group comparisons, in our specific experimental design, it is not well suited for this comparison. First, *limma* implicitly assumes that most of the analytes (genes / peptides / proteins) are not differentially expressed but this assumption is violated in the controlled spike-in datasets that we analyze, where a substantial fraction of the analytes is expected to be differentially expressed. Thus, it is not surprising that there are cases where *limma*’s differential analyses based on each of the peptide-level quantification tools apparently fails to control the FDR (Figure S2C). Notably, these apparent failures occur regardless of whether PIP is enabled as is evident by the estimated FDP in the no-PIP columns. Second, *limma* is based on t-tests that require at least two observations in each condition being compared. In practice, this can reward inaccurate implementations of PIP, because adding more observations enables a greater number of peptides/proteins to be quantified, regardless of the correctness of the additional observations. Thus a tool that erroneously adds more PIP identified peptides is rewarded by a *limma* differential expression test. Finally, MaxQuant had to be excluded from the *limma* analysis because MaxQuant does not output quantitative results for each run but instead averages all runs within an experimental condition. This type of output is incompatible with *limma*’s t-test based analysis.

To resolve these issues, we designed our own statistical evaluation to compare the effect of the choice of quantifications tool on the differential analysis. This evaluation takes advantage of the fact that for each of 10 pairwise abundance comparisons there is a known difference in the relative abundance between the respective spike-in levels. For example, when comparing the 1× and 3× spike-in conditions, the log_2_ fold-change is equal to log_2_(3*/*1) = 1.36. So in this case, all *E. coli* peptides/proteins should display a log_2_ fold-change of roughly 1.36, whereas all human peptides/proteins should display a log_2_ fold-change of about To determine the number of accurately quantified *E. coli* peptides/proteins, we count the number of those whose log_2_ fold-change falls within a distance of ∆*FC* of their expected fold-change (1.36 in our example). We define ∆*FC* so as to set the FDP at 5% by looking for its smallest value for which no more than 5% of the analytes whose log_2_ fold-change falls within the same window are human. This represents a 5% FDP, and is analogous to the adjusted p-value, or FDR cutoff, of 5% that is commonly used in *limma*. This approach is illustrated in Figure 4B.

**Figure 4:**
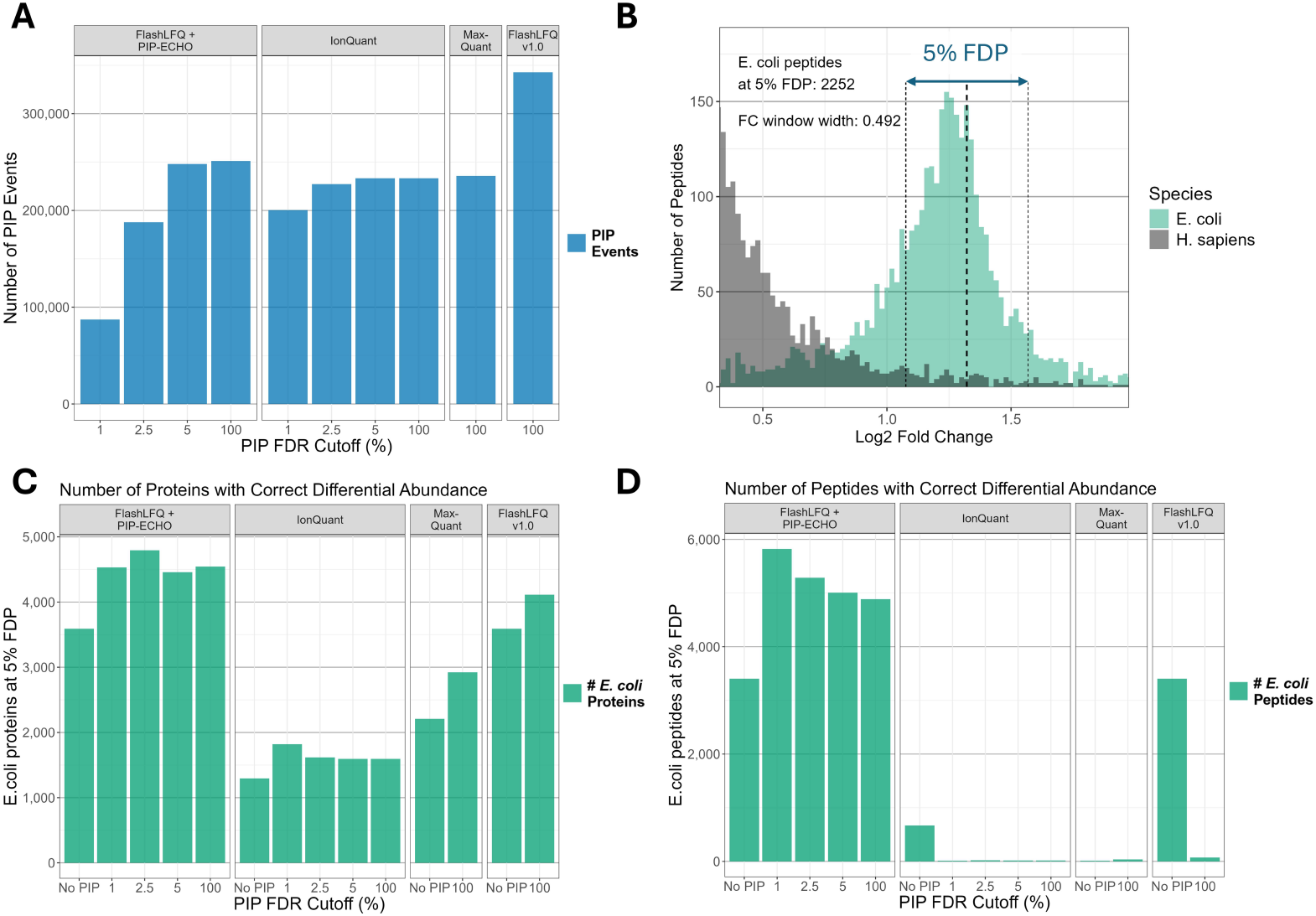
Differential Expression Analysis of an E.coli Spike-In Dataset. **A** Bar chart showing the number of PIP detections made by FlashLFQ v1.0, MaxQuant, IonQuant, and FlashLFQ v2.0 under a variety of analysis conditions. **B** Illustrated example of the sensitivity analysis. Results shown are from FlashLFQ v2.0, 5% PIP FDR, 1 ×vs ×2.5 spike-in comparison. **C & D** Sensitivity analysis of the different softwares. Plots show the number of differentially abundant peptides and proteins detected at 5% FDP (**C** - Protein level analysis, **D** - Peptide level analysis).

When we perform this differential abundance analysis, we find that at the same 5% FDP threshold, the analysis based on FlashLFQ+PIP-ECHO’s quantifications reports the largest number of differentially abundant proteins (4476 vs. 1819, 2921, 4112 for IonQuant, MaxQuant and v1.0 respectively) and peptides (5821 vs. 15, 39, and 73 for IonQuant, MaxQuant and v1.0 respectively). These results are shown in Figure 4C-D. This increase in performance occurs despite the fact that at 1% PIP FDR threshold, PIP-ECHO reports about half the number of PIPs that IonQuant and MaxQuant report and about a third of the number reported by FlashLFQ v1.0 (Figure 4A).

Figure 4 also shows that, for FlashLFQ v1.0 and IonQuant, enabling PIP actually decreases the number of differentially abundant peptides detected. This is because when PIP is performed, a greater number of human peptides are found (presumably falsely) to be differentially expressed between conditions. Notably, this is not the case for PIP-ECHO, where the addition of PIP transfers increases the number of peptides found to be differentially abundant. Interestingly, this observation holds regardless of PIP-ECHO’s selected FDR threshold, suggesting that PIP-ECHO is better at selecting correct transfers, thus generating cleaner quantitative data. This improved selection is probably thanks to a combination of the PEP-scoring of candidate traces with the competition between the randomized and properly predicted RT transfers.

## Discussion

While two-proteome experiments have been used in the past to evaluate PIP results, we show here for the first time how such experiments can be used to rigorously estimate the FDP of PIP, accounting for both peak-matching and peptide-identification errors. Our analysis suggests that only the new PIP-ECHO consistently controls the PIP FDR, particularly when considering single-cell analysis. We further demonstrate that quantifications based on combining PIP-ECHO with FlashLFQ enable a substantially more powerful differential abundance analysis compared with competing quantifications.

We plan on investigating future improvements to PIP-ECHO, including incorporating information from unidentified MS2 spectra, developing a better understanding of the effect of the FDR threshold that is used by the database-search tool to define potential donor peptides, and replacing the Percolator-like cross-validation scheme that is used in computing the PEP of candidate peak traces with the more rigorous Percolator-RESET type of approach [7]. We stress that PIP-ECHO was designed for DDA analysis, and it is not clear to us how easy it would be to extend it to DIA tools.

We note that PIP-ECHO controls the PIP FDR separately in each acceptor run, and PIPs from different acceptor runs that involve the same donor peptide are not independent. Thus, even if PIP-ECHO controls the FDR separately in each acceptor run, strictly speaking, it is not guaranteed to control the FDR in the combined list of PIPs from all runs. However, in practice with hundreds and even thousands of PIPs from each run, the law of large numbers implies that we can safely aggregate the PIPs across all runs while essentially controlling the FDR.

An implementation of PIP-ECHO is freely available through both FlashLFQ, a standalone tool for label-free quantification, and within the MetaMorpheus software suite. While the analysis of PIP-ECHO in this paper was within the framework of MetaMorpheus and FlashLFQ, in principle it can be combined with any database-search or LFQ tool.

## Methods

### Code and Data Availability

Code used for data analysis and figure generation can be found on GitHub: https://github.com/Alexander-Sol/ PIP-ECHOanalysis, https://doi.org/10.5281/zenodo.14112130. The Lim et al. two-proteome dataset can be found at the ProteomeXchange Consortium website with identifier PXD014415. The Shen et al. spike-in dataset can be found at the ProteomeXchange Consortium website with identifier PXD003881. The single-cell equivalent dataset can be found on PRIDE, with project identifier PXD037527. The *E. coli* dataset was generated specifically for this work and can be found at the ProteomeXchange Consortium website with the dataset identifier PXD057758, alongside the outputs from every software for every analysis that was performed.

FlashLFQ+PIP-ECHO was developed in the C# language as part of the mzLib library for mass spectrometry: https://github.com/smith-chem-wisc/mzLib. A standalone version of FlashLFQ+PIP-ECHO can be downloaded from https://github.com/smith-chem-wisc/FlashLFQ/releases.

### PIP-ECHO

The input to PIP-ECHO includes a list of potential donor peptides that are generated by searching a concatenated target-decoy database. In the results reported here we used MetaMorpheus to generate this list, but this can also be done using other search tools so long as the output contains both target and decoy PSMs. The number of peptides the tool reports can affect the power of PIP-ECHO, but as long as the tool is oblivious to the target/decoy label, PIP-ECHO should be able to control the PIP FDR.

Every peptide in the list of donor peptides is represented by one PSM — the highest scoring PSM that was matched to the donor peptide sequence. In addition to the highest scoring PSM, PIP-ECHO is also told the run from which the PSM originated, as well as a list of charges of all other PSMs from the same run that were matched to the same peptide sequence. PIP-ECHO then seeks to propagate the identity of the donor peptides to acceptor runs where the peptide was not MS2-detected.

PIP-ECHO attempts to match each potential donor peptide to two peak traces in each acceptor run. The first peak trace is sought within a window anchored at the peptide’s predicted acceptor RT, which is derived by locally matching the donor and acceptor runs’ RTs of peptides that were MS2-detected in both runs (Algorithm 5). The second peak trace in the same run is sought within a window anchored at a randomized RT drawn as follow. First, PIP-ECHO randomly draws a peptide from a list of MS2-detected peptides in the donor run that have: (a) an unambiguously associated peak trace (Algorithm 2), (b) a mass close to, but at least about 5 Da away from that of the original donor peptide, (c) a RT that is not too close to that of the original donor peptide, and (d) a stem or unmodified form that is distinct from that of the original donor peptide (Algorithm 13). Next, the RT of this randomly drawn donor peptide is mapped to the acceptor RT scale using the same Algorithm 5 that we used for mapping the RT of the original donor peptide.

Both the predicted-RT anchored window and the one anchored at the randomized RT have the same window size. Matching a donor peptide with a peak trace within either window requires scoring each eligible peak trace within the window and picking the highest scoring one. A peak trace is eligible if (a) it consists of one or more valid MS1 scans, i.e., the scans contain a peak that matches the theoretical most abundant isotopologue of the donor peptide, and they have a valid isotopic envelope matching the donor peptide’s theoretical envelope (Algorithm 7), and (b) the RT of the trace’s apex scan is within the window. An “apex scan” here is the MS1 scan in the peak trace with the maximal sum of log-intensities of peaks that match the theoretical isotopologues of the donor peptide (Algorithm 12). If neither the predicted-RT nor the randomized-RT based windows contain an eligible peak trace matching any of the observed donor peptide charges, then the window size is increased by 30 seconds. This process repeats until at least one such eligible peak trace is found or the maximal window size is reached and no peak trace is associated with the current donor (Algorithm 4).

Matches to peak traces are scored based on the following features: mass error, intensity, difference between observed and predicted/randomized retention time, isotopic distribution, and the number of MS1 scans in which an isotopic envelope was observed. Each feature is calibrated by computing the proportion of peak traces associated with MS2-detected peptides for which the value of the feature, as calculated for the match to the MS2-detected peptide, is at least as extreme as it is for the evaluated acceptor peak trace. For example, the mass error feature is the difference between the theoretical mass of the peptide’s most abundant ion and the deconvolved mass of the most abundant ion in the peak trace. This feature’s calibrated version is the proportion of MS2-detected peptides in the acceptor run for which the analogous difference, calculated with respect to their associated peak traces, is at least as large. The combined score for each matched peak trace is the geometric mean of these calibrated feature scores (Algorithm 11). This score is used to decide between multiple acceptor peak traces that are found within the same retention time window.

After peak trace matching has been performed for every acceptor run, the resulting list of candidate PIPs or matched peak traces are re-scored using a semi-supervised machine learning approach. This approach employs a gradient boosted binary decision tree classifier in order to assign every peak trace a posterior error probability (PEP) that, ideally, represents the likelihood that the peak trace is correctly matched. The same classifier, an implementation of the MART algorithm, is also used in MetaMorpheus, and a complete description is provided by Burges, 2010. [16, 2] Classification is performed in the style of Percolator using three-fold cross validation where the randomized-RT peaks are used as negative training examples and the predicted-RT peaks in the top 25% of all peaks are used as positive training examples. The first iteration of the semi-supervised learning model uses the combined score described in the preceding paragraph to select the top scoring peaks. Each subsequent iteration uses the PEP calculated in the previous iteration. Five rounds of training are performed in total, after which each peak trace has a well calibrated PEP (Algorithm 15).

Finally, for each donor peptide that is linked to both a predicted-RT and a randomized-RT acceptor peak trace, the peak trace with the higher PEP is removed. The remaining list of PIPs is sorted by increasing PEP and, given the desired FDR threshold *α*, the corresponding *k*(*α*) is computed using Equation (2). All predicted-RT target PIPs among the top *k*(*α*) PIPs are reported.

### Estimating False Discovery Proportion using Two-Proteome Datasets

#### Protein Databases

Each dataset was searched against a concatenated database composed of two separate databases. For the Lim et al. and single-cell equivalent datasets, the first database consisted of the *Saccharomyces cerevisiae* (UP000002311, 6060 proteins) proteome, whereas for the *E. coli* dataset, the database consisted of the *Escherichia coli* (UP000000625, 4404 proteins) proteome.

For all three datasets the second database was made of the human proteins fused with entrapment sequences designed to capture peptide identification errors. This database was created by appending an entrapment sequence to the end of each protein in the *Homo sapiens*(UP000005640, 82499 proteins) proteome. Entrapment sequences were created by randomly shuffling every residue in the target protein sequence while keeping all R and K residues (tryptic cleavage sites) in place. If any of the tryptic shuffled peptides were identical to a peptide found in the *Homo sapiens*, or in *Saccharomyces cerevisiae* (for the Lim et al. and single-cell datasets) or *Escherichia coli* (*E. coli* dataset) proteomes, they were removed. Half of each entrapment sequence was appended to the target protein sequence, resulting in a fused protein. This resulted in an approximately equal number of unique peptide sequences arising from the entrapment sequences and the *Homo sapiens* sequences.

All proteomes were dowloaded from UniProt in March of 2024. Additionally, the default contaminant database provided by each search engine (MaxQuant, MetaMorpheus, MSFragger) was used during search.

#### Data formatting

All data files were converted from .raw to .mzML format using msconvert prior to analysis by FragPipe+IonQuant and MetaMorpheus+FlashLFQ. MaxQuant analysis was performed using data in the .raw format because MaxQuant does not support the .mzML file type.

#### Database Search in MetaMorpheus

Each protein in each database was digested in silico to produce tryptic peptides. To be considered, peptides needed to have minimum length of 7, no more than two variable modifications, and no more than two missed cleavage sites. Oxidation of methionine was the only variable modification that was considered. For each peptide, a decoy peptide was generated by reversing every amino acid except for the C-terminus. In cases where this would produce an identical sequence, or a sequence that corresponds to a target peptide, a new decoy was generated by shuffling every amino acid except for lysine and arginine (in order to preserve tryptic cleavage sites). This procedure was repeated until a valid decoy was generated. If after five attempts no valid decoy could be generated, then the peptide was discarded from the database.

MetaMorpheus uses a gradient boosted decision tree classifier to assign a posterior error probability (PEP) to each PSM that denotes the probability that the spectrum was incorrectly matched to a peptide [16]. Initially, PSMs are assigned a MetaMorpheus score, which is 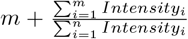 . Here, *m* is the number of fragment ions matched to the peptide, *n* is the total number of ions observed in the MS2 spectrum, and *Intensity*_*i*_ denotes the intensity of ion *i* within the MS2 spectrum.

Each peptide (peptidoform) is represented by the top-scoring PSM associated with that sequence. Peptides are arranged according to their MetaMorpheus scores, which are inherited from their top-scoring PSMs, and peptide-level q-values are calculated by leveraging the decoy PSMs as usually done in TDC. Specifically, let *T*_*k*_ be the number of target peptides among the top *k* scoring peptides, and let *D*_*k*_ be the number of decoy peptides among the same top *k* peptides. Then, the estimated FDR among that set of peptides is calculated as follows:

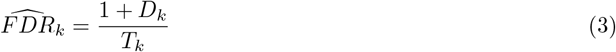

Q-values are calculated for each peptide by correcting the FDR estimates such that they increase monotonically as you move down the list. This is done by iterating backwards through the list, starting at *k* −1, and applying the following equation to each row:

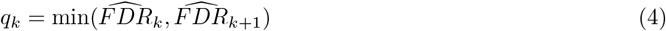

This procedure is also depicted in Algorithm 19.

All PSMs are split into four folds such that all peptides and PSMs derived from the same stem form (e.g., SAMPLER and SAM[Oxidation]PLER) are assigned to the same fold. Within each fold, the top-scoring PSM for each target peptide sequence with a peptide-level q-value *>*= 0.005 is labeled as a positive training example, and the top-scoring PSM for each decoy peptide sequence, regardless of its q-value, is labeled as a negative training example.

A classifier is trained on the positive and negative training examples from three out of the four folds, and this classifier is used to assign a PEP to every PSM in the held-out fold. This process is repeated until all four folds have been evaluated in this fashion.

Next, the PSMs are assigned so-called “(PSM) PEP q-values” by sorting them in increasing PEP score and applying the process described in Equations (3) and (4). Ties in the PEP score are broken by the MetaMorpheus score, then the mass error of the deconvolved precursor, then by scan number in this order.

A similar process is used to calculate peptide-level PEP q-values. Each peptide is represented by the most confident (lowest PEP) PSM associated with that peptide. Then, the peptides are arranged by PEP order, with MetaMorpheus score used as a tie-breaker. Finally, Equations (3) and (4) are again used to assign a PEP q-value to each peptide.

#### FlashLFQ v1.0

All PSMs were filtered to a PEP q-value of 0.01 before being passed into FlashLFQ v1.0 for quantification and PIP. Default argument were used for FlashLFQ.

#### FlashLFQ+PIP-ECHO

PSMs from MetaMorpheus were filtered to a PEP q-value of 0.01 before being passed into FlashLFQ+PIP-ECHO for quantification and PIP. Additionally, a list of peptides detected across all runs, along with their corresponding peptide-level PEP q-values, were passed as arguments to FlashLFQ+PIP-ECHO. Peptide-level PEP q-values were used to select donors for the PIP procedure. The donor peptide PEP q-value threshold differed depending on PIP FDR. PIP FDRs of 1%, 2.5%, 5% corresponds to donor peptide PEP q-value thresholds of 0.002, 0.005, and 0.01, respectively. For PIP FDRs of 100% (i.e., no PIP FDR threshold), a donor peptide PEP q-value threshold of 0.01 was used.

#### IonQuant

Datasets were searched and quantified using FragPipe v21.1, MSFragger v4.0, and IonQuant v1.10.12 [6, 11]. The “Default” workflow setting in FragPipe was used with several parameters changed as follows. The “Add decoys” option was enabled to add decoy and contaminant proteins to the protein database used for search. The “Run MS1 Quant” and “Match between runs (MBR)” options were enabled. The “MBR ion FDR” was adjusted based on the analysis being performed. For the spike-in dataset, the “Normalize intensity across runs” option was enabled. For the single-cell dataset, mixed samples were assigned to the “library” condition.

#### MaxQuant

Datasets were searched and quantified using MaxQuant v2.6.1.0 [3, 4, 23]. All default settings were used with the exception of the “Match between runs” option, which was enabled.

#### Data censoring

Data censoring was performed to estimate each PIP algorithm’s native peak matching error rate. This involved comparing peak traces that were unambiguously associated with MS2-detected peptides with corresponding peak traces identified by PIP using the same donor peptide but with the detecting MS2 spectra removed from the acceptor run. Data censoring was performed separately for each software tool that was evaluated as follows.

Let *MS*2 *Peptides*_*i*_ be the list of all peptides that were detected in the pure-sample run *i. MS*2 *Peptides*_*i*_ was filtered according to the following criteria:

a. The peptide must have been detected at an FDR *<* 0.1%
b. The peptide sequence must be derived from the *H*.*sapiens* proteome and must not occur in any other proteome used during the search.
c. The peptide must have been detected in at least one other run in the dataset with a score greater than the score of the peptide detection in run *i*.
d. The peptide must only be associated with one MS2 spectrum in run *i*.

Then, *Censored peptides*_*i*_ was created by randomly selecting 500 peptides from *MS*2 *Peptides*_*i*_ for censoring. Censoring was performed by editing the data file from run *i*. The MS2 spectrum associated with a peptide in *Censored peptides*_*i*_ was replaced with an MS2 spectrum that contained only one peak at 150 m/z. This was performed for every data file associated with a pure-sample run, and then the resulting censored data files were searched and quantified alongside the original mixed-sample data files. A different set of censored data files was generated for each software.

The results of the original and the censored data analyses were compared. Let *Native Peak Pairs*_*i*_ be the list of peak traces pairs where a peak trace corresponding to the same peptide was identified in the original as well as in the censored data analyses of run *i*.

Let *MS*2-*RT*_*x*,*i*_ be the retention time of the apex scan of the peak trace associated with *Peptide*_*x*_ in run *i* that was identified in the original analysis, and let *PIP* -*RT*_*x*,*i*_ be the corresponding retention time of the PIP involving *Peptide*_*x*_ in run *i* in the censored-data analysis. Let *PIP RT* -*Diff*_*x*,*i*_ = |*MS*2-*RT*_*x*,*i*_ −*PIP* -*RT*_*x*,*i*_| . Let *Native Peak Errors*_*i*_ be every pair in *Native Peak Pairs*_*i*_ where *PIP RT* -*Diff*_*x*,*i*_ *>* 1% the length of the LC gradient (Lim et al. - 54 seconds, *E. coli* - 36 seconds, Single-cell - 24 seconds). Then, let *All Native Peak* 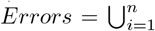 *Native Peak Errors*_*i*_ and *All Native Peak Pairs* 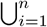 *Native Peak Pairs*_*i*_

#### Calculating FDP

1. Let *PIP* -*Identifications* be the list of all peak traces identified by PIP in pure (human-only) runs.
2. Let *Foreign MS*2-*Detections* be the list of all foreign species (yeast or E.coli, depending on the datasets) peptides that were detected in pure (human-only) runs.
3. All identifications in *PIP* -*Identifications* that correspond to a peptide in *Foreign MS*2-*Detections* were removed from *PIP* -*Identifications* (these are removed as suspected carryovers).
4. Each identification in *PIP* -*Identifications* was then assigned to one of the following lists:
  4.a. *Human PIP* -*IDs* - list of PIPs associated with *Homo sapiens* peptides,
  4.b. *Foreign PIP* -*IDs* - list of PIPs associated with a foreign species (*E. coli* or *S. cerevisaie*) peptide that is not present in the *Homo sapiens* proteome,
  4.c. *Entrapment PIP* -*IDs* - list PIPs associated with an entrapment peptide.
5. The estimated number of foreign peak matching errors is the observed number of foreign PIPs: *eFPE* = |*Foreign PIP* -*IDs*|.
6. The estimated number of peptide identification errors is derived from the entrapment PIPs: *ePIE* = |*Entrapment PIP* -*IDs*| *×S*, where, *S* is a scaling factor that corresponds to the ratio of the combined target database size to the entrapment database size: 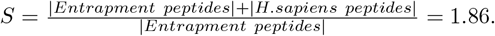.
7. The estimated number of native peak matching errors is

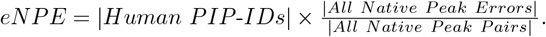
8. The estimated FDP is then equal to the sum of the three estimated error counts over the number of PIPs: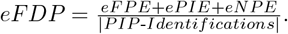.
9. Note the dividing each of the three estimated counts by |*PIP* -*Identifications*| gives an estimate of the FDP due to each of those three types of error.

### *E. coli* Two-Proteome Dataset

Human protein digest was obtained from Promega (MS Compatible Human Protein Extract, Digest, V695A) and resuspended in 5% acetonitrile, 0.2% formic acid buffer to a final concentration of 1 μg/μL. E. coli protein digest was obtained from Waters (MassPREP E. coli Digest Standard, 186003196) and resuspended in 5% acetonitrile, 0.2% formic acid buffer to a final concentration of 1 μg/μL. The mixed species sample was prepared by adding 3.5 μL of the E. coli protein digest solution to 35 μL of the human protein digest solution, for a final ratio of 10:1 human:E. coli peptides.

Both samples were analyzed over 10 replicate injections using an ultra-high performance LC-MS/MS (UPLC-MS/MS) system consisting of a Vanquish Neo ultra-high-pressure liquid chromatography system and an Orbitrap Fusion Lumos mass spectrometer (Thermo Fisher Scientific, RRID:SCR 020562). Injected peptide samples were loaded at a pressure of 400 bar onto a 25-cm long fused silica capillary nano-column packed with C18 resin (3.0-μm diameter, 130 Å pore size from Waters). Approximately 1 μg of peptide was loaded onto the column for each run. Peptides eluted over 140 min at a flow rate of 350 nL/min with the following gradient, where buffer A was aqueous 0.2% formic acid and buffer B was 80% acetonitrile with 0.2% formic acid: time 1 min-5% buffer B; time 35 min-30% buffer B; time 53 min-42% buffer B; time 60 min-55% acetonitrile; time 60.5– 85 min-85% buffer B. The nano-column was held at 60 C using a column heater constructed in-house.

The nanospray source voltage was set to 2,300 V. Full-mass profile scans were performed in the orbitrap between 375 and 1,500 m/z at a resolution of 120,000, followed by MS/MS higher-energy collisional dissociation (HCD) scans in the orbitrap of the highest intensity parent ions in a 3-s cycle time at 30% relative collision energy and 15,000 resolution, with a 2.5 m/z isolation window. Charge states 2–6 were included and dynamic exclusion was enabled with a repeat count of one over a duration of 30 s and a 10 ppm exclusion width both low and high. The automatic gain control (AGC) target was set to “standard,” maximum inject time was set to “auto,” and 1 μscan was collected for the MS/MS orbitrap HCD scan

### Differential Expression Analysis

Differential expression analysis was conducted using a spike-in dataset originally published by Shen et al.[15]. This dataset consists of *E. coli* and human lysate digests mixed at five different ratios, ranging from 3% to 9% total *E. coli* lysate (wt/wt). These ratios are denoted by the relative amounts of *E. coli* included in each: 1 ×, 1.5 ×, 2×, 2.5× and 3× . Four technical replicates were used for each concentration, resulting in 20 LC-MS/MS runs in total.

This dataset was searched against the *Escherichia coli* (UP000000625) and *Homo sapiens* (UP000005640) proteomes. Default contaminant databases for each database search engine were also included during search. Settings for each database search program are identical to those described in “Estimating False Discovery Proportion using Two-Proteome Datasets”. Quantification was performed using IonQuant, MaxQuant, FlashLFQ v1.0, and FlashLFQ+PIP-ECHO. For IonQuant and FlashLFQ+PIP-ECHO, PIP FDR thresholds ranging from 1-100% were tested. For all software, we compared quantification with and without PIP enabled. Normalization was performed using the normalization procedure implemented by each software.

The quantitative results for each file were analyzed as follows. The intensity values were log_2_ transformed, and missing or zero values were removed. Then, each of the five conditions were compa(r)ed (e.g., the 1× spike-in condition was compared to the 1.5×, 2×, 2.5× and 3× conditions), for a total of 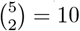 comparisons.

In each comparison, for each analyte (peptide or protein), the log_2_ intensity values that were reported for a given spike-in condition were averaged. Let *O*_*F C*_ denote the observed log_2_ fold change and let *E*_*F C*_ = log_2_(higher spike-in / lower spike-in) denote the expected log_2_ fold change of an *E. coli* peptide. For each analyte we calculated ∆_*F C*_ = *T*_*F C*_ − *O*_*F C*_, where perfectly accurate quantification should result in all *E. coli* analytes having a ∆_*F C*_ = 0, and all human peptides or proteins displaying a ∆_*F C*_ = *T*_*F C*_. Analytes were ordered by descending ∆_*F C*_, and the FDP among the top *k* analytes was calculated as 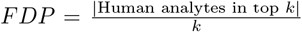 . Analytes that were ambiguous between the human and *E. coli* proteomes were excluded from the analysis. The number of *E. coli* analytes discovered at FDP = 0.05 were reported for each comparison and summed across all 10 comparisons.

Differential expression analysis using *limma* (v3.19) was conducted in a similar fashion[18]. Peptides/proteins with fewer than two observations in either condition were excluded from the analysis. Then, log_2_ transformed intensity values were used as an input into the standard *limma* workflow. A linear model was fit for each of the 10 pairwise comparisons using the `lmFit` function with default parameters and the `eBayes` function with “trend” set to True resulting in 10 sets of *limma* p-values, a p-value for each analyte. For each comparison the analytes with significant Benjamini-Hochberg-adjusted p-values at FDR thresholds of 0.01 and of 0.05 with a log_2_ fold change *>* 1.0 were deemed significantly differentially abundant. We reported the total number of *E. coli* peptides / proteins that were thus deemed significant summed up across the 10 comparisons, as well as the estimated FDP which we computed as the corresponding total number of human analytes that were deemed significant divided the sum of the human and *E. coli* totals.

## Acknowledgements

Figures 1, 2, and 3B were created with BioRender.com.

## Author contributions

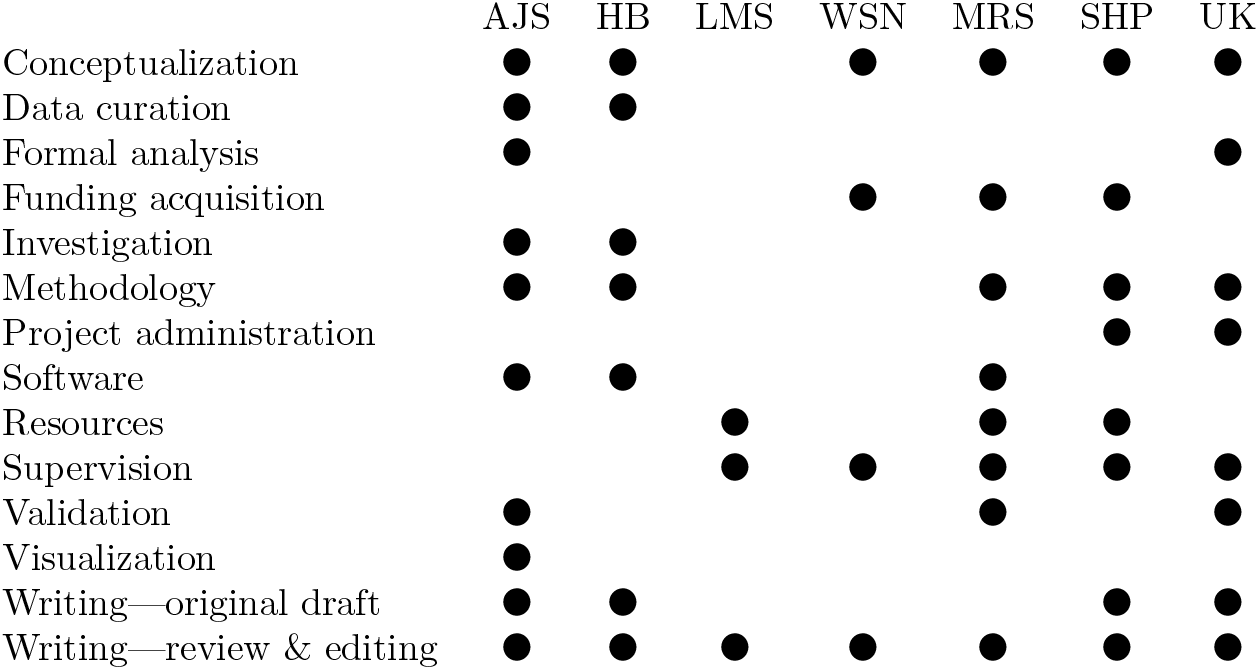

