## Supplemental for "Improved detection of differentially abundant proteins through FDR-control of peptide-identity-propagation"

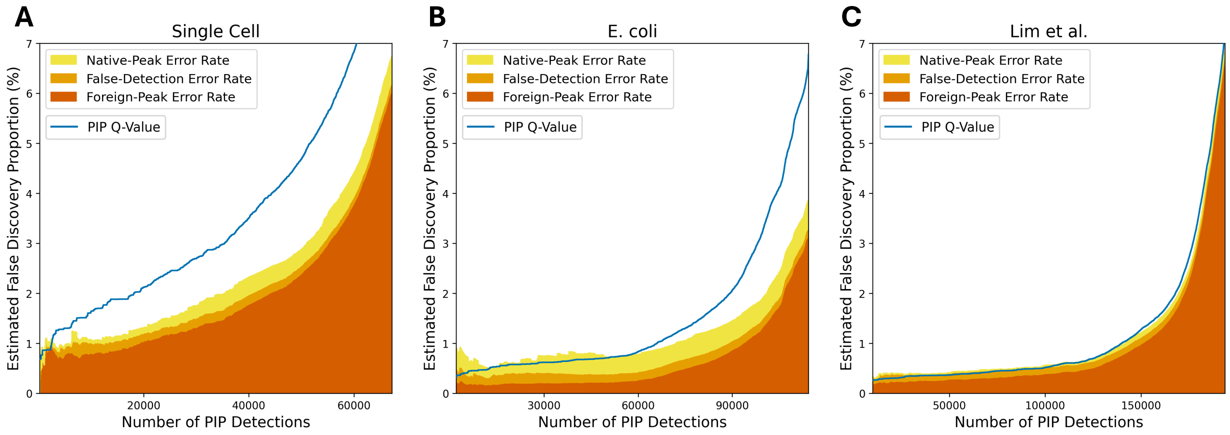


Figure S1: **Analysis of false discovery proportion versus false discovery rate** Stacked area plots showing the relationship between PIP q-values as calculated by PIP-ECHO and the estimates of the three different error rates that, when summed, give the estimated FDP . The PIP detections are ordered according to their PEP scores in decreasing significance (increasing PEP). **A** Analysis of a single-cell equivalent two-proteome dataset **B** Analysis of the *E. coli*+human dataset **C** Analysis of the Lim et al. dataset.

We find that PIP-ECHO's estimated FDR is mostly at or above the estimated FDP (some exceptions are visible for the smaller number of discoveries). The frequency of peptide-detection errors stays relatively constant for number of discoveries because we fixed the donor peptide FDR threshold at 0.002 (0.2%), and PIP-ECHO is unable to distinguish between accurate and inaccurate PSMs based on the properties of their associated peak traces. However, PIP-ECHO estimates the frequency of these errors and uses this information when calculating the estimated FDR for each set of PIP events. As the number of discoveries increases, peak-matching errors (particularly foreign-peak errors) become more common. This shows that PIP-ECHO is able to successfully differentiate accurate PIP transfers and erroneous ones. Taken together, these results show PIP-ECHO can reliably control the PIP FDR.

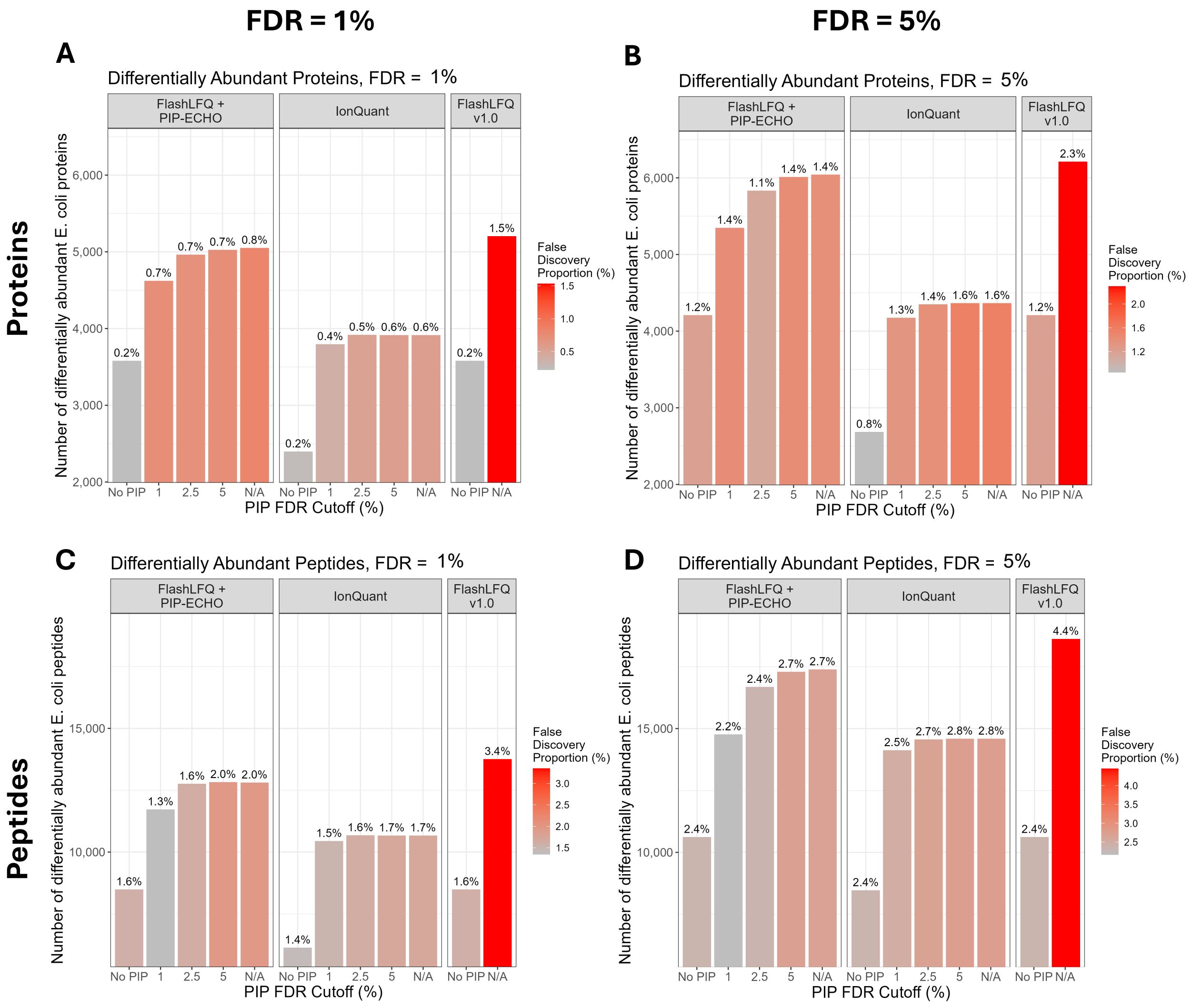


Figure S2: **Differential Expression Analysis using *limma*** Comparing *limma*'s DE analysis based on LFQ data generated by applying FlashLFQ+PIP-ECHO, IonQuant, and FlashLFQ v1.0 to a spike-in dataset originally published by Shen et al. [1]. This dataset consists of *E. coli* and human lysate digests mixed at 5 different ratios resulting in 20 LC-MS/MS runs in total. Pairwise analyses were performed between each of the 5 concentrations for a total of 10 *limma* pairwise DE analyses. Because this is a controlled dataset only *E. coli* peptides and proteins abundances are expected to change.

| Structure | Field | Description |
|---|---|---|
| $PSM$ | | A peptide-spectral-match obtained from a database search engine. Contains the following sub-fields: |
| $PSM$ | `seq` | the full sequence (including modifications) of the peptide associated with the PSM. |
| $PSM$ | `RT` | the retention time of the MS2 spectrum. |
| $PSM$ | `charge` | the charge of the precursor peptide that was isolated to generate the PSM. |
| $PSM$ | `score` | the score for the PSM. |
| $PSM$ | `decoy` | boolean indicating whether the peptide is a decoy or target. |
| $Tr$ | | An MS1 peak trace. A collection of isotopic envelopes, all of which are presumed to correspond to the same peptide precursor. Contains the following subfields: |
| $Tr$ | `PSM` | The PSM linked to the peak trace, temporally adjacent to at least one MS1 scan that contains an isotopic envelope. NULL for PIP peak traces. |
| $Tr$ | `donor` | For PIP peak traces, the donor peak trace that was used to identify the PIP peak trace. Mutually exclusive with `PSM`. |
| $Tr$ | `seq` | the full sequence (including modifications) of the peptide associated with the peak trace (and `PSM`, if `PSM` $\neq$ NULL). |
| $Tr$ | `run` | the identity of the LC-MS/MS data file, or run, in which the peak trace was found. |
| $Tr$ | `apex` | the MS1 scan that contains the most intense isotopic envelope (determined by summing the intensity of all observed isotopic peaks in each scan). |
| $Tr$ | `matchedPeak` | the peak in the apex scan that corresponds to the mass of the most abundant isotopologue of `seq`. |
| $Tr$ | `charges` | list of the precursor charges of all PSMs with the same peptide sequence. |
| $Tr$ | `score` | the PIP score, where higher scores are better. Only defined for for PIP peak traces. |
| $Tr$ | `scanCount` | the number of MS1 scans in which the isotopic envelope was observed. |
| $Tr$ | `ppmError` | the difference in ppm between the m/z of the most abundant (charged) isotope and its matched MS1 peak (the closest). |
| $Tr$ | `deltaRT` | the difference between predicted RT and the RT of the apex. |
| $Tr$ | `intensity` | log of the sum of the matched peak intensities in the apex scan, divided by the charge. |
| $Tr$ | `scanCount` | the number of scans containing valid isotopic envelope in peak trace. |
| $Tr$ | `corr` | the Pearson correlation between the theoretical isotopic envelope and the matched MS1 peaks in the apex scan. |
| $Tr$ | `PEP` | the PIP posterior error probability, where lower scores denote greater confidence in the PIP transfer. Only defined for for PIP peak traces. |
| $Tr$ | `Q` | the PIP Q-value. Only defined for for PIP peak traces. |
| $Tr$ | `isRealRT` | specifies whether or not the peak trace is associated with a predicted or random retention time. Only defined for for PIP peak traces. |
| $I_\pi$ | | The theoretical isotopic envelope of the peptide $\pi$. Contains the following sub-fields: |
| $I_\pi$ | `m` | a list of the masses of every isotopologue of $\pi$ with relative abundance $\geq 10\%$. |
| $I_\pi$ | `a` | a list of the relative abundances for every isotopologue of $\pi$ with relative abundance $\geq 10\%$. |

Table S1: **Classes and properties used in the algorithms below.**

| Structure/field | Description |
|---|---|
| *ASMP* | Apex Scan Match Parameters: acceptor-run-specific parameters associated with scoring the match between the apex scan and the charged peptide's theoretical isotopic envelope. The parameters are estimated from the MS2-detected peptides and their matched apex scans in the considered run. Contains the following sub-fields: |
| *ppmError.mean* | mean of the relative error (in ppm) between the m/z of the peptide's most abundant theoretical isotope and its matched ion peak in the apex scan; |
| *ppmError.SD* | corresponding SD. |
| *logIoC.mean* | mean of the log of the sum of theoretically-matched peak intensities divided by the charge state; |
| *logIoC.SD* | corresponding SD. |
| *oneMinusCor.shape* | estimated shape parameter (mean$^2$/variance) of the presumed gamma distribution of one minus the correlation between the intensities of the apex scan's matched peaks and their corresponding theoretical relative abundances.) |
| *oneMinusCor.rate* | corresponding estimated rate parameter (mean/variance) |
| *ARTMP* | Apex Retention Time Match Parameters: acceptor-donor-runs-specific parameters associated with scoring the agreement between the apex scan RT and the predicted RT. The parameters are estimated from peptides that are MS2-detected in both the donor and the acceptor runs. Contains the following sub-fields: |
| *RTdiff.mean* | mean of the difference between the acceptor peptide predicted and empirical (apex scan) RT; |
| *RTdiff.SD* | corresponding SD. |
| *PTMP* | Peak Trace Match Parameters: acceptor-run-specific parameters associated with scoring the match between the peak trace and the theoretical isotopic envelope. The parameters are estimated from the MS2-detected peptides and their matched peak traces in the considered run. Contains the following sub-fields: |
| *scanCount.mean* | mean number of valid isotopic envelope scans in trace peaks associated with the MS2-detected peptides in the acceptor run; |
| *scanCount.SD* | corresponding SD. |

Table S2: *peakParms* **- Parameters used in scoring the match between a precursor and an MS1 peak trace.**

| Parameter | Default | Description |
|---|---|---|
| $\Delta_A^{\max} RT$ | 1.5 minutes | maximum half-width of acceptor RT search window |
| $\Delta_m^{\min}$ | $5 \cdot 1.0078$ Da | minimum mass difference between candidate and donor |
| $\Delta_m^{\max}$ | $11 \cdot 1.0078$ Da | initial maximum difference |
| $\Delta_m^{\text{bound}}$ | $10^5$ Da | upper bound on maximum difference |
| $\Delta_{m/z}^{\text{bound}}$ | 10 ppm | upper bound on maximum m/z error in ppm |

Table S3: **Global parameters and their default values** .

**Algorithm 1:** `assignPeakTraceToPSM` - assign an MS1 peak trace to a reported PSM.

**Input:**

- $PSM$ a reported PSM with the following subfields:

**Output:**

- $Tr$ an MS1 peak trace associated with the PSM with the following subfields:

1. Find the maximal chain of scans so that
   - using $z = PSM.$`charge`, each scan has a peak at the m/z of the most abundant theoretical isotope of $PSM.$`seq`,
   - neighboring scans are separated by at most one MS1 scan,
   - $PSM.$`RT` is between the RTs of the two MS1 scans at the two ends of the chain.
2. If no such chain exists return an empty object. Otherwise, throw out of the chain scans that do not exhibit a valid isotopic envelope as determined by Algorithm 7 (potentially creating gaps more than one scan long).
3. Partition the chain into maximal sub-chains of scans for which each adjacent pair of scans are separated by at most one MS1 scan.
4. Further split sub-chains that are deemed to contain multiple local maxima so that each (sub) chain is essentially uni-modal (as detailed in the function `CutPeak`, line 1810 of the file `FlashLfqEngine.cs`).
5. If there exists a sub-chain whose RTs span across $PSM.$`RT` define it as the peak trace associated with $PSM$ and set $Tr.$`scanCount` and $Tr.$`PSM` accordingly. Otherwise, return an empty object.
6. Find the apex scan of the associated peak trace: the scan with the maximal sum of matched peak intensities and set $Tr.$`apex` and $Tr.$`matchedPeak` accordingly. (Algorithm 12).
7. Return the peak trace object $Tr$ with the above information.

**Algorithm 2:** `assignPeakTracesToDetectedPeptides` - assign an MS1 peak trace to each MS2-detected peptide for which it can be done unambiguously.

**Input:**

- $R$ list of LC-MS runs being analyzed with the following subfields:
  - `ID` the run's ID
  - `MS1` the MS1 data from the $k$th run in $R$
  - `PSMs` the run's reported PSMs

**Output:**

- $TR^2$ list of MS1 peak traces that were unambiguously associated with MS2-detected peptides from all runs with the following subfields:

$TR^2 \leftarrow \emptyset$; `// generated list of peak traces`<br>
**foreach** $r \in R$ **do**<br>
    $TR \leftarrow \emptyset$; `// run-specific raw list of peak traces`<br>
    **foreach** $PSM \in r.$*PSMs* **do**<br>
        $TR \leftarrow [TR,$ assignPeakTraceToPSM$(PSM, R)]$; `// add the PSM-associated MS1 peak trace to the list if it exists`<br>
    **end**<br>
    $\Pi \leftarrow$ `unique`$(TR[\bullet]$.`PSM.seq`$)$; `// all` $r$`'s MS2-detected peptides with an associate peak trace`<br>
    $TR_1 \leftarrow \emptyset$; `// keep only the run's best scoring trace for each peptide`<br>
    **foreach** $\pi \in \Pi$ **do**<br>
        $TR_\pi \leftarrow \{tr \in TR : tr.$`PSM.seq` $= \pi\}$; `// all the traces associated with` $\pi$<br>
        $k \leftarrow \text{argmax}\{TR_\pi[\bullet]$.`PSM.score`$\}$; `// trace with maximally scoring PSM`<br>
        $TR_\pi[k]$.`charges` $\leftarrow \{TR_\pi[\bullet]$.`PSM.charge`$\}$; `// record all charges associated with` $\pi$<br>
        $TR_\pi[k]$.`run` $\leftarrow r$.`ID`; `// add run ID`<br>
        $TR_1 \leftarrow [TR_1, TR_\pi[k]]$; `// add top scoring trace to this run's traces`<br>
    **end**<br>
    Remove from $TR_1$ any peak trace (characterized by the peak in its apex scan that is matched to the most abundant isotope) that appears multiple times in that list.<br>
    $TR^2 \leftarrow [TR^2, TR_1[\bullet]]$<br>
**end**

**Algorithm 3:** `computeMatchScoreParameters` - estimate the parameters of the distributions that are used to characterize and score the quality of a match between a candidate charged peptide and an MS1 peak trace.

**Input:**

- $Tr^2_A$ peak traces matched to MS2-detected peptides in runs $A$
- $\Pi^2_{AD}$ MS2-detected peptides common to both runs $A$ and $D$ ("MS2-common")
- $MS1_A$ run-$A$'s MS1 scans
- $MS1_D$ run-$D$'s MS1 scans

**Output:**

- $peakParms$ parameters required for scoring features of the match between the peptide and a peak trace (see Table S2 for details).

$mzErrors \leftarrow \{\texttt{mzError}(\pi) : \pi \in Tr^2_A\}$
$logIntensity \leftarrow \{\log_2 \texttt{intensity} \; |(Tr^2.A.\texttt{intensity} \in Tr^2_A\}$
$oneMinusCorr \leftarrow \{1 - \texttt{corr} \; |Tr.\texttt{corr} \in Tr^2_A\}$
$scanCounts \leftarrow \{\texttt{scanCount} \; |Tr.\texttt{scanCount} \in Tr^2_A\}$
$rtPredictionErrors \leftarrow \emptyset$
**foreach** $(Tr^2.A,\ Tr^2.D) \in Tr^2_{AD}$ **do**
    `// Calculate RT prediction errors for peak traces observed in both runs`
    $(RT^p_A, \Delta_A RT) \leftarrow \texttt{predictAcceptorRTWindow}(Tr^2.D.\texttt{apex.RT},\ TR^2_D,\ TR^2_A,\ \Pi^2_{AD},\ 1)$
    $predictionError \leftarrow RT^p_A - Tr^2.A.\texttt{apex.RT}$
    $rtPredictionErrors \leftarrow rtPredictionErrors \cup predictionError$
**end**
`// Populate peakParms`
$peakParms.ASMP.ppmError.mean \leftarrow \texttt{mean}(mzErrors)$
$peakParms.ASMP.ppmError.SD \leftarrow \texttt{sd}(mzErrors)$
$peakParms.ASMP.logIoC.mean \leftarrow \texttt{mean}(logIntensity)$
$peakParms.ASMP.logIoC.SD \leftarrow \texttt{sd}(logIntensity)$
$peakParms.ASMP.oneMinusCor.shape \leftarrow \texttt{mean}(oneMinusCorr)^2/\texttt{sd}(oneMinusCorr)^2$
$peakParms.ASMP.oneMinusCorr.rate \leftarrow \texttt{mean}(oneMinusCorr)/\texttt{sd}(oneMinusCorr)^2$
$peakParms.ARTMP.RTdiff.mean \leftarrow \texttt{mean}(rtPredictionErrors)$
$peakParms.ARTMP.RTdiff.SD \leftarrow \texttt{sd}(rtPredictionErrors)$
$peakParms.PTMP.scanCount.mean \leftarrow \texttt{mean}(scanCounts)$
$peakParms.PTMP.scanCount.SD \leftarrow \texttt{sd}(scanCounts)$
**return** $peakParms$

**Algorithm 4:** `generateCandidatePIPs` - generate a list of candidate PIPs by assigning acceptor peak traces to one donor peptide at a time. An assigned peak trace can lie within the donor's predicted run-$A$ RT window or in a random RT window (used as competition in FDR control).

**Input:**

- $DPT$ — potential donor peak traces
- $TR_A^2$ — run $A$ MS1 peak traces of MS2-detected peptides for which a peak trace could be assigned unambiguously
- $TR_D^2$ — same for run $D$
- $MS1_A$ — run-$A$'s MS1 scans
- *peakParms* — structure containing parameters used in scoring the match between a charged peptide and an MS1 peak trace (Table S2)

**Output:**

- $AP$ — list of all candidate PIPs using predicted as well as random RTs
  `isRealRT` — specifies whether or not the peak trace is associated with a predicted or random retention time.

$\Pi_{AD}^2 \leftarrow TR_A^2[\bullet]$.`PSM.seq` $\cap\, TR_D^2[\bullet]$.`PSM.seq`; `// MS2-detected peptides common to both runs` $A$ `and` $D$ `(''MS2-common'')`
$AP \leftarrow \emptyset$; `// a container for all PIPs`
**foreach** $Tr_d \in DPT$ **do**
  $\pi_d \leftarrow Tr_d$.`PSM.seq`; `// the donor candidate peptide with RT` $Tr_d$.apex.RT
  `/* look for acceptor peak traces matching` $\pi_d$ `using the predicted run-`$A$ `RTs of` $\pi_d$ `as well as of a randomly chosen peptide of similar mass */`
  $(RT_A^p, \Delta_A RT) \leftarrow$ `predictAcceptorRTWindow`($Tr_d$.apex.RT, $TR_D^2$, $TR_A^2$, $\Pi_{AD}^2$, $\Delta_A^{\max} RT$); `// center and half-width of predicted run-`$A$ `RT window of` $\pi_d$ `(Algorithm 5)`
  $\tau_{\pi_d}^r \leftarrow$ `drawRandomRT`($\pi_d$, $Tr_d$.apex.RT, $TR_D^2$, $\Delta_A RT, \Pi_D^2$); `// predicted RT of randomly selected peptide (Algorithm 13)`
  $(RT_A^r, \_) \leftarrow$ `predictAcceptorRTWindow`($\tau_{\pi_d}^r$, $TR_D^2$, $TR_A^2$, $\Pi_{AD}^2$, $\Delta_A^{\max} RT$); `// map the RT to run` $A$ `(Algorithm 5)`
  **repeat**
    **if** *notNull*$(RT_A^p)$ **then**
      `// look for a` $\pi_d$`-matching peak trace in run` $A$ `using` $\pi_d$`'s predicted RT (Algorithm 6)`
      $(Tr, S) \leftarrow$ `matchDonorPeptide`($Tr_d$, $RT_A^p$, $\Delta_A RT$, $MS1_A$, *peakParms*);
    **end**
    **if** *notNull*$(RT_A^r)$ **then**
      `// again but now using a randomized RT`
      $(Tr^r, S^r) \leftarrow$ `matchDonorPeptide`($\pi_d$, $RT_A^r$, $\Delta_A RT$, $MS1_A$, *peakParms*);
    **end**
    $\Delta_A RT \leftarrow \Delta_A RT + 0.25$; `// increase the window size by 1/2 minute`
    **if** $S > -\infty \,||\, S^r > -\infty$ **then**
      **if** $S > -\infty$ **then**
        `// add the candidate PIP structure tagging it as a real RT)`
        $AP \leftarrow [AP, (\text{seq} = \pi_d, \text{peakTrace} = Tr, \text{score} = S, \text{isRealRT} = \text{TRUE})]$;
      **end**
      **if** $S^r > -\infty$ **then**
        `// add the candidate PIP structure tagging it as a random-RT)`
        $AP \leftarrow [AP, (\text{seq} = \pi_d, \text{peakTrace} = Tr^r, \text{score} = S^r, \text{isRealRT} = \text{FALSE})]$;
      **end**
      **break**; `// found at least one acceptor peak trace so no need to extend the RT window`
    **end**
  **until** $\Delta_A RT > \Delta_A^{\max} RT$;;
**end**
**return** $Ap$

**Algorithm 5:** `predictAcceptorRTWindow` - returns the initial acceptor RT window for matching a donor peptide RT

**Input:**

- $\tau_{\pi_d}$ the donor peptide's presumed retention time (RT)
- $TR_A^2$ run $A$ MS1 peak traces of MS2-detected peptides for which a peak trace could be assigned unambiguously
- $TR_D^2$ same for run $D$
- $\Pi_{AD}^2$ MS2-detected peptides common to both runs $A$ and $D$ ("MS2-common")
- $\Delta_A^{\max} RT$ maximum half-width of acceptor RT search window

**Output:**

- $(RT_A^p, \Delta_A RT)$ center of acceptor window and its half-width

`// Step 1: Map` $\tau_{\pi_d}$ `to Run` $A$

`// First find MS2-common donor peptides distinct from` $\pi_d$ `with donor run RT` $(RT(\pi, TR_D^2))$ `up to 1/2 minute away from it`

$\Pi_{AD_0} \leftarrow \{\pi \in \Pi_{AD}^2 \,:\, 0 < |RT(\pi, TR_D^2) - \tau_{\pi_d}| \leq 0.5\}$;

`//` $\Pi_{AD}^c$ `defined next contains the 3 closest MS2-common peptides that elute within 0.5 minute on either side of` $\pi_d$

$\Pi_{AD}^c \leftarrow \{\pi \in \Pi_{AD_0} \,:\,$ one of three smallest values with $RT(\pi, TR_D^2) - \tau_{\pi_d} \geq 0\}$;

$\Pi_{AD}^c \leftarrow \Pi_{AD}^c \cup \{\pi \in \Pi_{AD_0} \,:\,$ one of three largest values with $RT(\pi, TR_D^2) - \tau_{\pi_d} \leq 0\}$;

$\Delta_{AD} RT \leftarrow \{RT(\pi, TR_A^2) - RT(\pi, TR_D^2) \,:\, \pi \in \Pi_{AD}^c\}$; `// corresponding 6` $A \rightarrow D$ `RT shifts`

**if** $\Delta_{AD} RT == \emptyset$ **then**

  **return** $(\texttt{NULL}, -1)$; `// cannot reliably map` $\tau_{\pi_d}$ `run` $A$

**end**

$RT_A^p \leftarrow \tau_{\pi_d} + \text{median}(\Delta_{AD} RT)$; `// run A's RT of` $\pi_d$ `is predicted using local shifts`

`// Step 2: Determine` $\Delta_A RT$`, the maximal allowed run-`$A$ `RT distance to` $RT_A^p$

**if** $|\Delta_{AD} RT| > 1$ **then**

  $\Delta_A RT \leftarrow \min\{6 \cdot \sigma(\Delta_{AD} RT), \Delta RT_{max}\} \;/\; 2$; `// window of considered retention time range is` $RT_A^p \pm 3\times$ `the standard deviation`

**end**

**else**

  $\Delta_A RT = \Delta_A^{\max} RT$; `// cannot reliably estimate the window size`

**end**

**return** $(RT_A^p, \Delta_A RT)$

**Algorithm 6:** `matchDonorPeptide` - looks to match a donor peptide to a peak trace within the acceptor window

**Input:**

- $Tr_d$ — the donor peak trace
- $RT_A^p$ — center of acceptor window
- $\Delta_A RT$ — half-width of acceptor window
- $MS1_A$ — run-$A$'s MS1 scans
- $peakParms$ — structure containing parameters used in scoring the match between a charged peptide and an MS1 peak trace (Table S2)

**Output:**

- $(Tr_{\max}, S_{\max})$ — the optimal matching run-$A$ MS1 peak trace and its score

$\pi_d \leftarrow Tr_d$.`seq`; `// the donor candidate peptide`

$I_{\pi_d} \leftarrow$ `commonIsotopes`$(\pi_d)$; `//` $(I_\pi[k].m, I_\pi[k].\mathtt{a}) =$ `theoretical (mass, relative abundance) of` $k$`th isotopologue of` $\pi$ `with` $I_\pi[k].\mathtt{a} \geq 10\%$

$i_{\max} \leftarrow \mathrm{argmax}(I_{\pi_d}[\bullet].\mathrm{a})$; `// most abundant isotope of` $\pi_d$

**foreach** $z \in Tr_d.charges$ **do**

  `// iterate through the set of all charges associated with donor run PSMs with` $\pi_d$

  $mz_{i_{\max}} \leftarrow I_{\pi_d}[i_{\max}].m/z +$ `mass`(proton); `// corresponding` $m/z$ `of` $i_{\max}$

  $MS1_A^c \leftarrow \{\sigma \in MS1_A : |\sigma.RT - RT_A^p| \leq \Delta_A RT$ && `isPeakInScan`$(mz_{i_{\max}}, \sigma, \Delta_{m/z}^{\text{bound}})\}$;

    `// all run-A MS1 scans within the considered RT window with an ion peak within` $\Delta_{m/z}^{\text{bound}}$ `(default 10) ppm of` $mz_{i_{\max}}$

  $MS1_A^c \leftarrow \{\sigma \in MS1_A^c :$ `isValidIsotopicEnvelope`$(\sigma, I_{\pi_d})\}$; `// further retain only scans that present a valid isotopic envelope (Algorithm 7)`

  $Tr_A^c \leftarrow$ `getCandidateTraces`$(MS1_A^c)$; `// partition the valid MS1 scans into peak traces (Algorithm 21)`

  $Tr_{\max} \leftarrow \emptyset$; $S_{\max} \leftarrow -\infty$; `// maximally matching peak trace and its score`

  **foreach** $Tr \in Tr_A^c$ **do**

    **if** $Tr \neq \emptyset$ **then**

      $S \leftarrow$ `scoreTraceMatch`$(Tr, I_{\pi_d}, RT_A^p, peakParms)$; `// score the match to the considered peak trace (Algorithm 11)`

      **if** $S > S_{\max}$ **then**

        $S_{\max} \leftarrow S$; $Tr_{\max} \leftarrow Tr$;

      **end**

    **end**

  **end**

**end**

**return** $(Tr_{\max}, S_{\max})$;

**Algorithm 7:** `isValidIsotopicEnvelope` - returns TRUE iff the MS1 scan contains a valid isotopic envelop

**Input:**
- $\sigma$ - the MS1 scan represented as a structured array:
  - $\sigma[k]$.`mz` - the m/z of the $k$th peak of $\sigma$
  - $\sigma[k]$.`i` - the intensity of the peak
- $I_\pi$ - the theoretical isotopic envelope of the peptide $\pi$:
- $z$ - the charge state of the analyzed peptide precursor

**Output:** TRUE iff the $\sigma$ is deemed to present a valid envelope

$\Delta M_I \leftarrow \texttt{mass}(^{13}\text{C}) - \texttt{mass}(^{12}\text{C})$; `// the basic isotopic mass shift`
$I_\pi \leftarrow [(\min(I_\pi[\bullet].\texttt{m}) - \Delta M_I, 0), I_\pi]$; `// Adding a null isotopologue to account for a potential off-by-one error`
$I^c_\pi \leftarrow \texttt{calibrateEnvelope}(I_\pi, \sigma, z)$; `// calibrate the isotopes mass on the mass shift of the most abundant isotope (Algorithm 8)`
$\rho_0 \leftarrow \texttt{getIsotopicEnvelopeCor}(\sigma, I^c_\pi, z)$; `// find the correlation between the MS1 scan and the theoretical isotopic envelope (Algorithm 9)`
**if** $\rho_0 == \emptyset$ **then**
  **return** *FALSE*; `// an envelope requires at least two isotopes with MS1 peaks`
**end**
$I^-_\pi[\bullet].\texttt{a} \leftarrow I^c_\pi[\bullet].\texttt{a}$;
$I^-_\pi[\bullet].\texttt{m} \leftarrow I^c_\pi[\bullet].\texttt{m} - \Delta M_I$;
$\rho_{-1} \leftarrow \texttt{getIsotopicEnvelopeCor}(\sigma, I^-_\pi, z)$; `// find the correlation between the MS1 scan and the theoretical envelope shifted by the isotopic mass shift`
**if** $\rho_{-1} == \emptyset$ **then**
  $\rho_{-1} \leftarrow -1$;
**end**
$I^-_\pi[\bullet].\texttt{m} \leftarrow I^c_\pi[\bullet].\texttt{m} + \Delta M_I$;
$\rho_{+1} \leftarrow \texttt{getIsotopicEnvelopeCor}(\sigma, I^-_\pi, z)$; `// theoretical envelope shifted by the isotopic mass shift in the other direction`
**if** $\rho_{+1} == \emptyset$ **then**
  $\rho_{+1} \leftarrow -1$;
**end**
$isValid \leftarrow \rho_0 > 0.7 \;\&\&\; \rho_{-1} - \rho_0 < 0.1 \;\&\&\; \rho_{+1} - \rho_0 < 0.1$; `// comparisons with` $\rho_{\pm 1}$ `are to prevent an off-by-one error`
**return** $isValid$

**Algorithm 8:** `calibrateEnvelope` - returns the theoretical isotopic envelope calibrated on the mass shift of the most abundant isotope

**Input:** Same as Algorithm 7
**Output:** $\rho$ - the Pearson correlation

$MXA \leftarrow \text{argmax}\, I_\pi[\bullet].\texttt{a}$; `// most abundant isotopologue`
$MSP \leftarrow \text{argmin}\, |I_\pi[MXA].\texttt{m} - z \cdot \sigma[\bullet].\texttt{mz}|$; `// closest MS1 peak`
$\Delta m \leftarrow I_\pi[MXA].\texttt{m} - z \cdot \sigma[MSP].\texttt{mz}$; `// mass shift of the most abundant isotope`
$I^c_\pi.\texttt{a} \leftarrow I_\pi.\texttt{a}$;
$I^c_\pi[\bullet].\texttt{m} \leftarrow I_\pi[\bullet].\texttt{m} - \Delta m + z \cdot \texttt{mass}(^1\text{H})$; `// calibrating the mass of the charged isotope on that mass shift`
**return** $(I^c_\pi, 10^6 \cdot (-\Delta_m + z \cdot mass(^1H))/I_\pi[MXA].m)$; `// (calibrated envelope, calibrating mass shift (error) in ppm)`

**Algorithm 9:** `getIsotopicEnvelopeCor` - returns the Pearson correlation between the MS1 scan and the theoretical isotopic envelope

**Input:** Same as Algorithm 7
**Output:** $\rho$ - the Pearson correlation

$MP \leftarrow$ `findMatchedPeaks`$(\sigma, I_\pi, z)$; `// get list of matched theoretical and MS1 peaks (Algorithm 10)`
**if** *length*$(MP) < 2$ **then**
  **return** $\emptyset$; `// need at least two matched peaks`
**end**
`// find the correlation between the relative abundance of the theoretical isotopologue and the intensity of its matched MS1 peak`
$\rho \leftarrow$ `computePearsonCorrelation`$(MP[\bullet].\texttt{a}, MP[\bullet].\texttt{i})$;
**return** $\rho$;

**Algorithm 10:** `findMatchedPeaks` - returns a list of matched theoretical and MS1 peaks

**Input:** Same as Algorithm 7
**Output:** $MP$ - list of matched theoretical and MS1 peaks

$MP \leftarrow \emptyset$; **foreach** $(m, a) \in I_\pi$ **do**
  `// look for a matching MS1 peak for each considered theoretical isotopologue`
  ; $k \leftarrow \texttt{argmin}\{|\sigma[\bullet].\texttt{mz} - (m/z))|\}$; `// find the closest MS1 peak`
  **if** $|\sigma[k].mz - (m/z))/(m/z)| < \Delta_{m/z}^{bound} \cdot 10^{-6}$ **then**
    $MP \leftarrow [MP, (a, \sigma[k].\texttt{i})]$; `//` $m/z$ `should be no more` $\Delta_{m/z}^{\texttt{bound}}$ `ppm away`
  **end**
**end**

**Algorithm 11:** `scoreTraceMatch` - score the match between the candidate peak trace and the theoretical isotopic envelope

**Input:**

| | |
|---|---|
| • $Tr$ | the peak trace |
| • $I_\pi$ | the theoretical isotopic envelope of the peptide $\pi$: |
| • $z$ | the charge state of the analyzed peptide precursor |
| • $RT_A^p$ | predicted acceptor run RT |
| • $peakParms$ | structure containing parameters used in scoring the match between a charged peptide and an MS1 peak trace (Table S2) |

**Output:**

| | |
|---|---|
| • matchQ | structure evaluating the match quality with the following sub-fields: |
| `peakError` | the difference in ppm between the m/z of the most abundant (charged) isotope and its matched MS1 peak (the closest) |
| `S.peakError` | corresponding score |
| `deltaRT` | the difference between predicted and empirical RT |
| `S.deltaRT` | corresponding score |
| `intensity` | log of sum of matched apex peak intensities divided by the charge |
| `S.intensity` | corresponding score |
| `scanCount` | the number of valid isotopic envelope scans in peak trace |
| `S.scanCount` | corresponding score |
| `corr` | the correlation between the theoretical isotopic envelope and the matched MS1 apex peaks |
| `S.corr` | corresponding score |
| `combined` | the geometric mean of the above 5 feature scores |

`// First, find the apex scan, sum of its matched peaks intensities, theoretical envelope calibrated on the apex scan and the corresponding mass shift in ppm (Algorithm 12)`

$(\sigma_a, SaI, I_\pi^c, peakError) \leftarrow$ `findApexScan`$(Tr, I_\pi^c)$;

$p \leftarrow peakParms$; `// shorter name`

$matchQ$.`S.peakError` $\leftarrow 2 \cdot \Phi\left(-\frac{|peakError - p.ppmError.mean|}{p.ppmError.SD}\right)$; `// mass ppm error score (`$\Phi$` is the `$N(0,1)$` CDF)`

$\Delta RT \leftarrow RT_A^p - \sigma_a.RT$; `// difference between predicted and empirical RT`

$matchQ$.`S.deltaRT` $\leftarrow 2 \cdot \Phi\left(-\frac{|\Delta RT - p.RTdiff.mean|}{p.RTdiff.SD}\right)$; `// `$\Delta_{RT}$` score`

$logIoC \leftarrow \log_2(SaI/z)$; `// intensity gauge`

$matchQ$.`S.intensity` $\leftarrow 2 \cdot \Phi\left(-\frac{|logIoC - p.logIoC.mean|}{p.logIoC.SD}\right)$; `// intensity score`

$scanCount \leftarrow |Tr|$; `// number of valid isotopic envelope scans in trace`

$matchQ$.`S.scanCount` $\leftarrow 2 \cdot \Phi\left(-\frac{|scanCount - p.scanCount.mean|}{p.scanCount.SD}\right)$; `// scan count score`

$\rho \leftarrow$ `getIsotopicEnvelopeCor`$(\sigma_a, I_\pi^c, z)$; `// find the correlation between the MS1 scan and the theoretical isotopic envelope (Algorithm 9)`

$matchQ$.`S.corr` $\leftarrow F_{\Gamma[p.oneMinusCor.shape, p.oneMinusCor.rate]}(1-\rho)$; `// the isotopic correlation score where `$F_{\Gamma[\alpha,\lambda]}$` is the Gamma CDF with shape `$\alpha$` and rate `$\lambda$

$matchScore \leftarrow 100 \cdot (matchQ$.`S.peakError` $\cdot\, matchQ$.`S.deltaRT` $\cdot\, matchQ$.`S.intensity` $\cdot\, matchQ$.`S.scanCount` $\cdot\, matchQ$.`S.corr`$)^{1/5}$; `// geometric mean of the features scores`

**return** $matchScore$

**Algorithm 12:** `findApexScan` - returns the apex scan: the scan with maximal sum of matched peak intensities

**Input:**

- $Tr$ the peak trace
- $I_\pi$ the theoretical isotopic envelope of the peptide $\pi$:
- $z$ the charge state of the analyzed peptide precursor

**Output:**

- $\sigma_a$ the apex scan
- $SaI$ sum of its matched peaks intensities
- $I^c_\pi$ isotopic envelope calibrated on mass shift of the most abundant isotope
- $\Delta m$ ppm mass difference between the most abundant ion and intense peak

$\sigma_a \leftarrow \emptyset$; $SaI \leftarrow -\infty$ `// apex scan and sum of its matched peaks intensities`
**foreach** $\sigma \in Tr$ **do**
  $(I^c_\pi, \Delta m) \leftarrow$ `calibrateEnvelope`$(I_\pi, \sigma, z)$; `// calibrate the isotopes mass on the mass shift of the most abundant isotope (Algorithm 8)`
  $MP \leftarrow$ `findMatchedPeaks`$(I^c_\pi, \sigma, z)$; `// find the matching peaks`
  **if** $\sum MP[\bullet].i > SaI$ **then**
    $\sigma_a \leftarrow \sigma$; $SaI \leftarrow \sum MP[\bullet]$.i; `// update candidate apex`
  **end**
**end**
**return** ($\sigma_a$, $SaI$, $I^c_\pi$, $\Delta m$)

**Algorithm 13:** `drawRandomRT` - draw a random RT for the donor peptide

**Input:**

- $\pi_d$ the donor peptide
- $\tau_{\pi_d}$ its presumed retention time (RT)
- $TR^2_D$ run $D$ MS1 peak traces of MS2-detected peptides for which a peak trace could be assigned unambiguously
- $\Delta_A RT$ half-width of acceptor window
- $\Pi^2_D$ MS2-detected peptides in the donor run

**Output:**

- $\tau^r_{\pi_d}$ random RT for $\pi_d$

`// The random RT is defined as the RT of a randomly-selected MS2-detected peptide from the donor run. This peptide is drawn from a list of candidates,` $\Pi^c$`, that is sequentially refined (see Table S3 for definitions of` $\Delta^{\min}_m, \Delta^{\max}_m, \Delta^{\text{bound}}_m$`).`
**while** $\Delta^{\max}_m \leq \Delta^{bound}_m$ **do**
  $\Pi^c \leftarrow \{\pi \in \Pi^2_D$ : $|$`mass`(`mostAbundantIsotope`$(\pi)$) $-$ `mass`(`mostAbundantIsotope`$(\pi_d)$)$| \in [\Delta^{\min}_m, \Delta^{\max}_m]\}$; `// MS2-detected donor run peptides with the prescribed mass difference to` $\pi_d$`.`
  $\Pi^c \leftarrow \{\pi \in \Pi^c : |RT(\pi, TR^2_D) - \tau_{\pi_d}| > 4 \cdot \Delta_A RT\}$; `// RT of candidate peptide cannot be too close to` $\pi_d$`'s`
  $\Pi^c \leftarrow \{\pi \in \Pi^c$ : `removePTMs`$(\pi) \neq$ `removePTMs`$(\pi_d)\}$; `// the ''stem form'' or unmodified versions of the candidate must differ from` $\pi_d$`'s`
  **if** $\Pi^c \neq \emptyset$ **then**
    **return** `randomlyDrawFromSet`$(RT(\Pi^c, TR^2_D)$); `// randomly draw the run-`$D$ `RT of one of the candidates`
  **end**
  **else**
    $\Delta^{\max}_m \leftarrow 10 \cdot \Delta^{\max}_m$
  **end**
**end**
**return** *NULL* `// failed to define a random RT`

**Algorithm 14:** `PIP-ECHO` - the main procedure

**Input:**

- $TR^2$ — list of MS1 peak traces that were unambiguously associated with MS2 detections from all runs
- $R$ — list of LC-MS runs being analyzed with the following subfields:
  - `ID` — the run's ID
  - `MS1` — the MS1 data from the $k$th run in $R$
  - `PSMs` — the run's reported PSMs
- $peakParms$ — structure containing parameters used in scoring the match between a charged peptide and an MS1 peak trace (Table S2)
- $\alpha$ — The desired FDR of the PIP procedure

**Output:**

- $AP$ — A list of All PIPs reported by PIP-ECHO

$S \leftarrow$ `unique`$(TR^2[\bullet]$.`seq`); `// list of distinct PIP donor peptide sequences`
$DPT \leftarrow \emptyset$; `// list of donor peak traces for PIP`
**foreach** $s \in S$ **do**
  `// For each donor peptide` $s$ `find its highest scoring MS2 detection and select its associated peak trace for PIP`
  $k \leftarrow \text{argmax}_{k=1,\ldots,\texttt{length}(TR^2)}\{TR^2[k].\texttt{PSM.score} : TR^2[k].\texttt{seq} == s\}$;
  $DPT \leftarrow [DPT, TR^2[k]]$;
**end**
$ACP \leftarrow \emptyset$ `// list of All Candidate PIPs`
**foreach** $r \in R$ **do**
  `// allocate to each run its MS1 peak traces in` $TR^2$ `(unambiguously assigned MS2-detections)`
  $r.\texttt{TR2} \leftarrow \{TR^2[k] : TR^2[k].\texttt{run} == r.\texttt{ID}\}$;
**end**
`// Iterate through each potential acceptor run`
**foreach** $r_a \in R$ **do**
  $ACP_a \leftarrow \emptyset$ `// list of All Candidate PIPs for acceptor run` $r_a$
  $DPT_a \leftarrow \{Tr \in DPT : Tr.\texttt{seq} \notin r_a.\Pi^2\}$; `// PIP must involve a peptide that was not MS2-detected in run A`
  `// Iterate through each potential donor run`
  **foreach** $r_d \in R \setminus r_a$ **do**
    $DPT_{ad} \leftarrow \{Tr \in DPT_a : Tr.\texttt{run} == r_d.\texttt{ID}\}$; `//` $r_d$`'s potential donor peak traces`
    $ACP_a \leftarrow ACP_a \cup$ `generateCandidatePIPs`$(DPT_{ad}$, $r_a$.`TR2`, $r_d$.`TR2`, $r_a$.`MS1`, $peakParms)$
  **end**
  $ACP_a \leftarrow$ `resolvePIPsAmbiguities`$(ACP_a$, $r_a$.`TR2`); `// acceptor peak traces should be unambiguously assigned (Algorithm 22)`
  $ACP_a \leftarrow \{$`addField`$(CP$, `run`, $r_a$.`ID`$) : CP \in ACP_a\}$; `// add the run ID to each PIP`
  $ACP \leftarrow ACP \cup ACP_a$
**end**
$ACP \leftarrow$ `PEPscoring`$(ACP)$; `// rescore the PIPs using multiple features (Algorithm 15)`
`// Iterate through each run, calculating the Q value for each PIP peak trace`
**foreach** $r \in R$ **do**
  $AP_r \leftarrow \{PIP \in ACP : PIP.\texttt{run} == r.\texttt{ID}\}$
  $AP_r \leftarrow \text{calculateFDR}(AP_r)$
**end**
$AP \leftarrow \{PIP \in ACP : PIP.Q \leq \alpha\}$
**return** $AP$

**Algorithm 15:** `PEPscoring` - calculate PEP

**Input:**

- $AP$ A list of All PIPs reported by PIP-ECHO

**Output:**

- $AP$ PIPs after PEP scoring

Sort $AP$ in descending score order<br>
$i \leftarrow \left\lfloor \frac{|AP|}{4} \right\rfloor$ `// Calculate the index for the 25th percentile`<br>
$S_t \leftarrow AP[i]$.`score` `// Calculate PIP score threshold that determines which peak traces will be used as positive training examples`<br>
$G \leftarrow \emptyset$ `// Initialize list to hold groups of peak traces and group all acceptor peak traces that are derived from the same donor`<br>
**foreach** $d \in AP[\bullet]$.*`donor`* **do**<br>
    $g \leftarrow \{Tr \in AP | Tr.$`donor` $== d\}$<br>
    $G \leftarrow [G,\ g]$<br>
**end**<br>
$P \leftarrow$ `partitionGroups`$(G)$ `// Split` $G$ `into three partitions,` $P_1$`,` $P_2$`, and` $P_3$<br>
`// Define the sets of PIPs that will be used to train the classifier`<br>
$Ex^+ \leftarrow \emptyset$ `// Positive training examples`<br>
$Ex^- \leftarrow \emptyset$ `// Negative training examples`<br>
**for** $i \leftarrow 1$ **to** *3* **do**<br>
    $Ex^+[i],\ Ex^-[i] \leftarrow$ `getTrainingExamples`$(P_i,\ S_t,\ null)$<br>
**end**<br>
Swap groups between partitions such that each partition has an equal number of positive and negative training examples. This is done using the EqualizeDonorGroupIndices function, line 234 of PepAnalysisEngine.cs<br>
$P \leftarrow$ `crossValidationScoring`$(\{P_1,\ P_2,\ P_3\},\ Ex^+,\ Ex^-)$ `// Calculate a PEP score for every PIP peak trace`<br>
`// Iterative training`<br>
**for** $i \leftarrow 1$ **to** *4* **do**<br>
    $Ex^+ \leftarrow \emptyset$<br>
    $Ex^- \leftarrow \emptyset$<br>
    **for** $j \leftarrow 1$ **to** *3* **do**<br>
        $A_j \leftarrow \emptyset$ `// Collect all PIPs in partition` $j$<br>
        **foreach** $g \in P_j$ **do**<br>
            $A_j \leftarrow A_j \cup \{Tr \in g\}$<br>
        **end**<br>
        Sort $A_j$ in ascending PEP order<br>
        $k \leftarrow \left\lfloor \frac{|A_j|}{4} \right\rfloor$ `// Calculate the index for the 25th percentile`<br>
        $PEP_t \leftarrow A_j[i]$.`PEP` `// Calculate PEP threshold that determines which peak traces will be used as positive training examples`<br>
        $Ex^+[j],\ Ex^-[j] \leftarrow$ `getTrainingExamples`$(P_j,\ null,\ PEP_t)$<br>
    **end**<br>
    $P \leftarrow$ `crossValidationScoring`$(\{P_1,\ P_2,\ P_3\}, Ex^+,\ Ex^-)$<br>
**end**<br>
**return** *`AP`* `// Return all PIPs with calculated PEPs`

**Algorithm 16:** `partitionGroups`

**Input:**

- $G$  a list of PIP groups

**Output:**

- $P$  A list of three partitions, each containing one third of the groups in $G$

$i \leftarrow 0$<br>
**while** $i < |G|$ **do**<br>
    $n \leftarrow (i \bmod 3) + 1$<br>
    $P_n \leftarrow P_n \cup G[i]$<br>
    $i \leftarrow i + 1$<br>
**end**<br>
**return** $\{P_1,\ P_2,\ P_3\}$

**Algorithm 17:** `getTrainingExamples`

**Input:**

- $G$  a list of groups containing all acceptor PIPs from a single donor peptide
  - $G[k]$.`AP`  acceptor peak traces in the $k^{th}$ group in $G$
- $S_T$  PIP score threshold that determines which PIPs traces should be used as positive training examples
- $PEP_T$  PEP threshold that determines which PIPs should be used as positive training examples

**Output:**

- $Ex^+$  a list of PIPs to be used as positive training examples
- $Ex^-$  a list of PIPs to be used as negative training examples

`// Instantiate empty lists to hold the positive and negative training examples`<br>
$Ex^+ \leftarrow \emptyset$<br>
$Ex^- \leftarrow \emptyset$<br>
**foreach** $g \in G$ **do**<br>
    $Ex^- \leftarrow Ex^- \cup \{Tr \in g\ |Tr.$`isRealRT == False`$\}$ `// Add random-RT PIPs to the negative training examples`<br>
    **if** $S_T \neq null$ **then**<br>
        `// Add predicted-RT PIPs above` $S_t$ `to the positive training examples`<br>
        $Ex^+ \leftarrow Ex^+ \cup \{Tr \in g\ |Tr.$`isRealRT == True AND` $Tr.$score $\geq S_T\}$<br>
    **end**<br>
    **else if** $PEP_T \neq null$ **then**<br>
        `// Add predicted-RT PIPs below` $PEP_t$ `to the positive training examples`<br>
        $Ex^+ \leftarrow Ex^+ \cup \{Tr \in g\ |Tr.$`isRealRT == True AND` $Tr.$score $\leq PEP_T\}$<br>
    **end**<br>
**end**<br>
**return** $(Ex^+,\ Ex^-)$

**Algorithm 18:** `crossValidationScoring`

**Input:**

- $P$ a list of partitions containing groups of peak traces
  $P[k]$.G all groups of peak traces in the $k^{th}$ partition of P
- $Ex^+$ A list of positive training examples from each partition
- $Ex^-$ A list of negative training examples from each partition

**Output:**

- $P$ partions where every peak trace has an updated PEP score

```
// Cross-validation classification
for i ← 1 to 3 do
    b = (i + 1) % 3
    c = (i + 2) % 3
    // Combine training examples from partitions b and c
    Ex+ = Ex+[b] ∪ Ex+[c]
    Ex- = Ex-[b] ∪ Ex-[c]
    TrainedModel ← A gradient boosted binary decision tree trained on Ex+ and Ex-
    foreach g ∈ P_i do
        foreach Tr ∈ g do
            Tr.PEP ← TrainedModel.Predict(Tr)
        end
    end
end
return P /* A list of partitions where every PIP has been assigned a PEP */
```

**Algorithm 19:** `calculateFDR` - calculate FDR for all PIP peak traces in a given acceptor run

**Input:**

- $A_r$ list of all PIP peak traces from acceptor run $r$

**Output:**

- $A_r$ PIP peak traces after Q-value calculation

```
/* Order PIP peaks according to their PEP */
Sort the elements of A_r in monotone increasing PEP order
foreach Tr_i ∈ A_r do
    /* Calculate an intermediate Q-Value for Tr_i by counting the number of different
       types of PIP peak traces in {Tr_n|n ≤ i} */
    // number of target-peptide peak traces with predicted retention times
    T_pr ← |{Tr^p ∈ Tr_n|Tr.Donor.PSM.Decoy == False}
    // number of target-peptide peak traces with random retention times
    T_r ← |{Tr^r ∈ Tr_n|Tr.Donor.PSM.Decoy == False}
    // number of decoy-peptide peak traces with predicted retention times
    D_pr ← |{Tr^p ∈ Tr_n|Tr.Donor.PSM.Decoy == True}
    // number of decoy-peptide peak traces with random retention times
    D_r ← |{Tr^r ∈ Tr_n|Tr.Donor.PSM.Decoy == True}

    Q_i ← computeQValue(T_pr, T_r, D_pr, D_r)
end
/* Correct intermediate Q-values to ensure they are in monotone increasing order */
for i = |Q| − 1 to 1 do
    Q_i ← min(Q_i, Q_{i+1})
end
return Q /* Return a list of every Q-value */
```

**Algorithm 20:** `computeQValue`

**Input:**
- $T_{pr}$ - number of target-peptide PIP peak traces with a predicted retention time
- $T_r$ - number of target-peptide PIP peak traces with a random retention time
- $D_{pr}$ - number of decoy-peptide PIP peak traces with a predicted retention time
- $D_r$ - number of decoy-peptide PIP peak traces with a random retention time

**Output:**
- $Q$ A Q-value

```
/* Estimate the number of errors arising from peptide-detection errors */
```
$\epsilon_d \leftarrow max(0, D_{pr} - D_r)$
```
/* Calculate the hybrid error Q-value */
```
$Q \leftarrow \frac{1+T_r+\epsilon_d}{T_{pr}}$

**return** $Q$

**Algorithm 21:** `getCandidateTraces` - partition the valid MS1 scans into candidate peak traces.

**Input:**
- $MS1^c_A$ - list of valid MS1 scans

**Output:** $Q$ - A Q-value $Tr^c_A 1$ - list of candidate peak traces

`// Step 1: partition the list of valid MS1 scans` $MS1^c_A$ `into maximal chains of scans with each adjacent pair of scans separated by at most one MS1 scan (a chain can include a single valid scan).`

```
// Step 2: further split chains that are deemed to contain multiple local maxima
   so that each (sub) chain is essentially uni-modal (as detailed in the function
   CutPeak, line 1810 of the file FlashLfqEngine.cs).
// Step 3: return the list of MS1 chains, each corresponding to one candidate peak
   trace characterized by the peak in its apex scan that is matched to the most
   abundant isotope.
```

**Algorithm 22:** `resolvePIPsAmbiguities` - remove PIPs that score lower than another PIP that shares the same acceptor peak trace, as well as those whose peak trace is assigned to an MS2-detected acceptor peptide.

**Input:**
- $Ap$ initial list of all candidate PIPs
- $\Pi^2_A$ MS2-detected peptides in runs $A$

**Output:**
- $Ap_1$ updated filtered list of all candidate PIPs

```
// Step 1: an acceptor peak trace, as characterized by the peak in its apex scan
   that is matched to the most abundant isotope, can only be used in a single PIP
// Step 2: an acceptor peak trace cannot already be assigned to an MS2-detected
   acceptor peptide
```

**Algorithm 23:** `equalizePartitions` - Swap groups of PIPs between partitions to equalize the number of positive training examples (predicted retention time, high PIP score) and negative training examples (random retention time) in each partition

**Input:**

- $P$ initial list of three partitions
  $P[k].G$ The list of groups associated with partition $k$
- $Ex^+$ A list of positive training examples from each partition
- $Ex^-$ A list of negative training examples from each partition

**Output:**

- $P$ updated partitions
- $Ex^+$ updated positive training examples from each partition
- $Ex^-$ updated negative training examples from each partition

`// This algorithm summarizes the behaviour of the EqualizeDonorGroupIndices function, which can be found at line 234 of PepAnalysisEngine.cs, within mzLib/FlashLFQ`

`// Step 1: Compare` $|Ex^+[1]|$ `to` $|Ex^+[2]|$ `and` $|Ex^-[1]|$ `to` $|Ex^-[2]|$ `to determine the difference in the number of positive and negative training examples between partitions 1 and 2`

`// Step 2: Iterate through groups in` $P[1]$ `and` $P[2]$ `and attempt to swap groups such that the difference in the number of training examples is reduced.`

`// Step 3: Repeat steps 1 and 2, but this time consider` $P[2]$ `and` $P[3]$ `and` $|Ex^\pm[2]|$ `and` $|Ex^\pm[3]|$

`// Step 4: Repeat steps 1-3 for a total of three times`

**return** $(P,\ Ex^+,\ Ex^-)$ `// Return balanced partitions and training sets`
